# Compositional computations in the primate frontal cortex

**DOI:** 10.64898/2026.11.05.764642

**Authors:** Cheng Tang, Mehrdad Jazayeri

## Abstract

A hallmark of intelligence is the ability to apply familiar operations to new inputs without relearning. How neural circuits carry a learned computation into states never visited during training remains unclear. Here we combined neural recordings, circuit modeling and human experiments to investigate compositional generalization. Monkeys learned to combine a start location with a displacement specified by the direction and duration of an optic-flow stimulus, then encountered novel combinations of these familiar inputs. Frontal population activity separated the computation into spatial and temporal components: start location and flow direction established the neural state, from which activity evolved along shared timing dynamics. Novel combinations elicited accurate responses when trajectories maintained this organization, whereas departures predicted systematic behavioral biases. Circuit models revealed a three-way alignment problem: input representations, recurrent dynamics and readout operations must jointly implement the required computation. Mismatches produced systematic biases that increased with distance from trained conditions. Humans exhibited similar biases when familiar sensory cues and spatial transformations were linked through an unfamiliar mapping. Together, these findings show how the coordination of reusable neural resources supports new computations, and how its disruption produces predictable failures of generalization.

## Introduction

A hallmark of intelligence is the ability to apply familiar operations to unfamiliar inputs. Having learned to add, we can sum quantities we have never combined before; having learned to mentally rotate a shape, we can rotate new shapes. In each case, a learned function is applied to configurations never previously experienced. This capacity to use known functions to compute new results allows finite knowledge to support open-ended behavior. Yet how neural circuits carry a learned computation beyond the combinations on which they were trained remains poorly understood.

Recent work has found that neural populations and artificial networks encode task variables in structured representations ^1–8^ that support abstract knowledge ^9–12^, working memory ^13^, memory scaffolds ^15^ and cognitive maps ^5,14,16,21,22^. These representations allow downstream circuits to access information about individual variables across changes in other variables and context. Yet preserving information about individual variables does not explain how a circuit can transform and combine them correctly in configurations never encountered during training.

Addressing this gap requires understanding how representations and dynamics work together. For a computation implemented through neural dynamics to generalize, those dynamics must perform the appropriate transformation when novel input combinations place the circuit in states not encountered during training. The central question is how latent dynamics are sculpted in regions of state space never visited during training ^23^.

To investigate this question, we trained monkeys to combine a start location with a displacement conveyed by an optic-flow stimulus. The task required integrating flow direction over time and adding the resulting displacement to the start location. Training covered the individual inputs but only a subset of their combinations, allowing us to probe the learned dynamics with novel pairings. We recorded population activity in the dorsomedial frontal cortex (DMFC) to ask whether those dynamics preserved the organization needed for accurate computation beyond the trained combinations.

Animals generalized successfully to some combinations but made systematic errors on others. Neural recordings revealed that start location and flow direction set the neural state, from which activity evolved along a shared temporal dimension. When novel combinations preserved this organization, animals composed the inputs accurately; when trajectories departed from it, behavior showed predictable errors. Recurrent neural network models showed how alignment between input representations, recurrent dynamics and downstream operations could sustain the computation in untrained regions of state space, whereas mismatches produced graded failures. Complementary human experiments revealed similar failures when familiar inputs and operations were linked through a novel mapping. Together, these findings suggest that compositional generalization depends on whether neural dynamics preserve the structure of a learned computation beyond the states encountered during training.

## Results

### Vector composition task

We trained two monkeys to compute a target location by combining a Start location with a displacement (Fig. 1a). Each trial began with central fixation and presentation of the Start at a point on a 7×7 grid. A coherent-motion stimulus (Flow) was then presented within a circular aperture centered on fixation. The direction and duration of the Flow specified the direction and magnitude of the displacement, respectively. After a brief delay, the animal reported the target location with a saccade. Denoting the Start vector by **s**, the displacement unit vector by **d**, and Flow duration by t, the correct vector composition was **s** + **d**t. Juice reward decreased linearly with the distance between the reported and correct endpoints, reaching zero at an error of half the displacement magnitude (Fig. 1b). The task included 48 Start locations, four cardinal Flow directions, and integer displacement magnitudes that kept the endpoint on the grid and off the fixation point, yielding 564 valid conditions.

**Figure 1.**
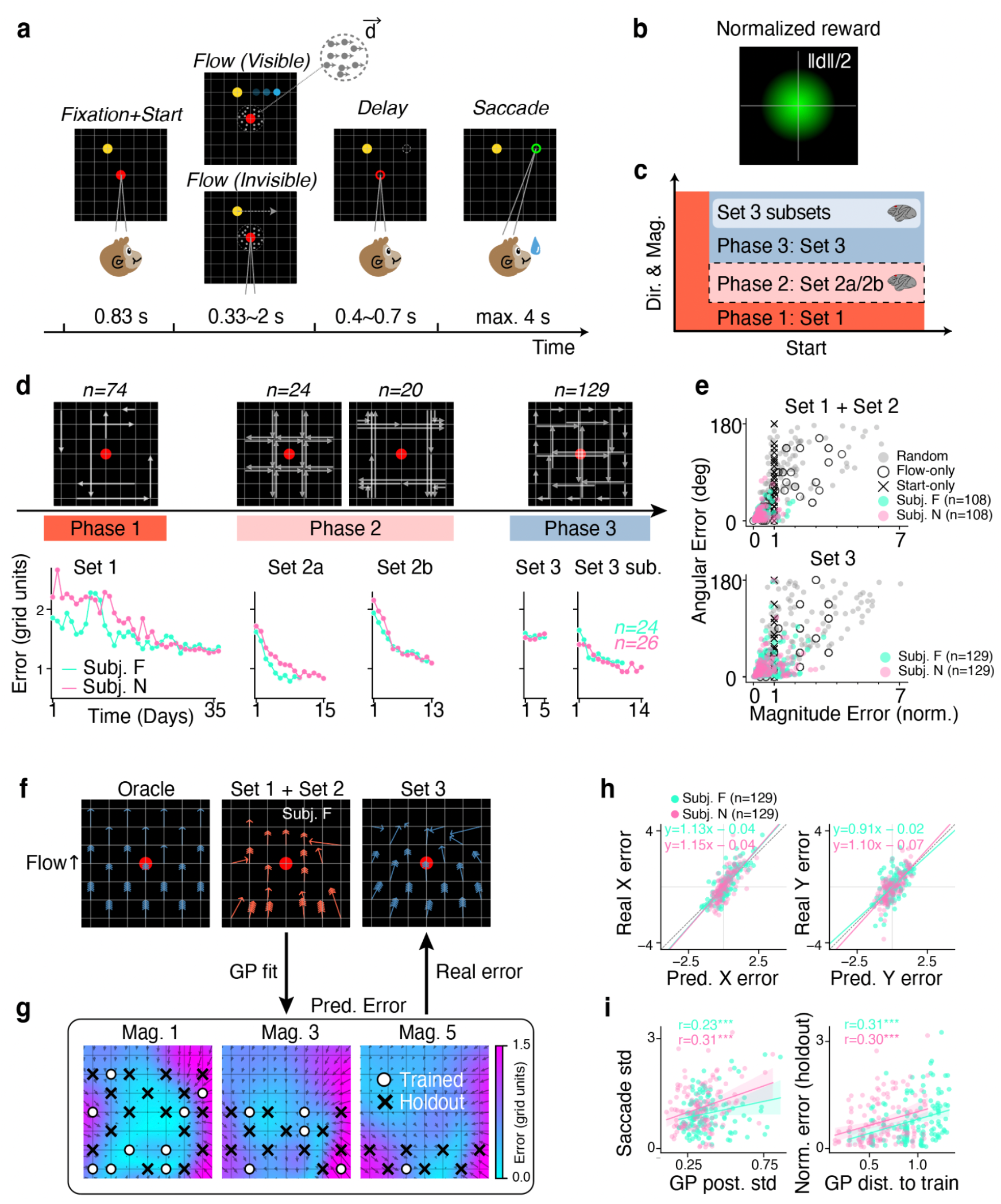
Compositional behavior in a vector-addition saccade task. **a**, Trial structure. The subjects (F and N) fixated at the center of a 7×7 grid while a Start location (yellow dot) appeared at a grid intersection. A coherent motion Flow stimulus specified a displacement vector through its direction and duration. In visible trials (top), a blue dot traced the displacement; in invisible trials (bottom), no visual guidance was provided. Monkeys reported the inferred target location via saccade after a variable delay. Trial timing is indicated below. **b**, Reward schedule. Juice reward scaled linearly with response accuracy, reaching maximum for correct responses and zero when error exceeded half the displacement magnitude (ǁdǁ/2). **c**, Experimental design. Three phases: Phase 1 (Set 1, orange), initial training; Phase 2 (Set 2, dashed box), neural characterization; Phase 3 (Set 3, blue), compositional generalization. Brain icons indicate phases with neural recordings. **d**, Behavioral performance across sessions. Top: example conditions for each phase (*n*, number of task conditions). Bottom: mean response error (grid units) across sessions for both animals (Subj. F, cyan; Subj. N, pink). **e**, Magnitude error (normalized by flow magnitude) versus angular error for different strategies. Each point represents one condition. Strategies: Random (uniform sampling), Flow-only (add Flow to origin, ignoring Start), Start-only (saccade to Start, ignoring Flow), and actual animal behavior. Top, Set 1 + Set 2 conditions; bottom, Set 3 conditions. *n*, number of conditions. **f**, Mean saccade responses for upward Flow conditions (Subj. F). Each arrow points from the Start to the saccade location, with length normalized by the displacement magnitude (correct saccades all normalize to unit length). The number of caps on each arrow indicates the actual Flow duration. Left, oracle (perfect performance). Middle, Set 1 + Set 2 (trained with visible conditions). Right, Set 3 (generalization). **g**, Gaussian process (GP) error field fit to the last five sessions of Set 1 and Set 2. Colormap, predicted error magnitude; arrows, predicted error direction. GP predicts an error map for each direction × magnitude combination; only upward Flow of magnitudes 1, 3 and 5 is shown. Circles, trained (Set 1 + Set 2) conditions; crosses, Set 3 conditions. **h**, GP-predicted (fit on Set 1 and Set 2) versus actual error for x (left) and y (right) axes across Set 3 holdout conditions. Each point is one Set 3 condition. **i**, Proximity effect. Each point is one Set 3 condition. Left, GP-predicted (fit on Set 1 and Set 2) posterior uncertainty versus trial-to-trial saccadic variability. Right, GP kernel distance of a Set 3 condition to the most accurate same-direction Set 1 + Set 2 condition versus behavioral error of the Set 3 condition. ***, p < 0.001.

We organized the experiment into three phases (Fig. 1c,d and Figs. S1.1 and S1.2). In Phase 1, we trained animals on a subset of conditions to establish the basic task contingencies. In Phase 2, we introduced two additional subsets and performed large-scale recordings in DMFC to characterize the neural representations and dynamics supporting task performance. In Phase 3, we tested novel Start–Flow combinations and examined the behavioral and neural signatures of successful generalization and systematic errors. Below, we first describe behavior across all three phases, then examine neural activity in Phases 2 and 3.

### Behavior – Phase 1

In Phase 1, we trained animals on 74 Start–Flow conditions (Set 1), selected to include all possible values of **s**, **d**, and t, but only a subset of their combinations. To facilitate learning, we initially presented a moving visual cue that traced the displacement from the Start during the Flow period. The cue faded gradually as performance improved (Methods). Both animals reached asymptotic performance on the training set (Fig. 1d; 35 sessions for Subj. F and Subj. N).

### Behavior – Phase 2

In Phase 2, we introduced two structured subsets, Set 2a and Set 2b, on alternating days. Set 2a varied Flow direction at a fixed duration across Start locations (24 conditions), whereas Set 2b varied Flow duration (magnitudes 1, 3 and 6) at a fixed direction for each Start location (20 conditions). Ten of these conditions also appeared in Set 1, so Sets 1 and 2 together comprised 108 unique training conditions. Animals were trained on each set before neural recordings began. To maintain a baseline reward rate, previously learned Set 1 conditions were randomly interleaved on 25% of trials. Performance stabilized within a few sessions (Fig. 1d; Set 2a: 7 sessions for Subj. F, 10 for Subj. N; Set 2b: 7 sessions for both animals).

To assess whether animals used both Start and Flow, we decomposed their responses into normalized magnitude error and angular error and compared them with three model observers: a Random observer that sampled response locations uniformly, a Start-only observer that reported the Start location, and a Flow-only observer that applied the displacement from the grid origin. The animals’ errors clustered closer to zero on both measures than those of any model observer, indicating that their responses incorporated both Start and Flow information (Fig. 1e).

To visualize behavior for individual conditions, we represented each animal’s mean response as a vector from the Start to the mean saccade endpoint. We normalized its length by the required displacement magnitude, so that an accurate response had unit length, and used one arrowhead per grid unit to indicate the required displacement.

Under this convention, perfect responses (oracle) appear as unit vectors pointing in the correct direction, angular errors appear as rotations, and magnitude errors produce vectors shorter or longer than unit length (Fig. 1f). This visualization revealed both accurate performance and systematic biases. For example, for upward Flow, responses from Start locations on either side of the grid deviated toward the center (Fig. 1f). We later examine how these behavioral biases relate to departures from the neural dynamics supporting accurate composition.

### Behavior – Phase 3

In Phase 3, we tested generalization to 129 novel Start–Flow pairings (Set 3) without visual guidance. We limited this initial probe to five sessions to minimize additional learning. Performance changed little over these sessions, with mean error decreasing by 3.8% in Subj. F and 0.2% in Subj. N. We then selected a subset of conditions for neural recordings in each animal, including both consistently accurate responses and systematic biases (Fig. S1.3). These subsets allowed us to examine the neural signatures of successful and unsuccessful generalization (Fig. 1d).

Applying the same observer-model analysis to Set 3 revealed a pattern similar to that in the trained conditions (Set 1 + Set 2). Both animals’ magnitude and angular errors remained closer to zero than those of the Random, Flow-only, and Start-only observers (Fig. 1e), indicating that their responses incorporated Start location and both Flow direction and magnitude even in novel combinations.

Examining individual conditions revealed that Set 3 errors resembled those in the trained sets. For the upward Flow, animals responded accurately in many conditions, but responses from Start locations on either side of the grid again deviated toward the center (Fig. 1f).

The similarity between trained and novel conditions suggested that behavioral errors might vary smoothly across task space, allowing errors in novel combinations to be predicted from those in trained conditions. To test this possibility, we fitted a Gaussian process (GP) to errors in the last five sessions of Set 2a and Set 2b (Set 1 conditions interleaved) as a smooth function of Start location and Flow, and used it to predict errors in the held-out Set 3 conditions (Fig. 1g; Methods).

GP predictions were strongly correlated with observed errors in the held-out conditions along both saccade axes (x: Subj. F, r = 0.83, p = 1.7 × 10 ⁻ ³ ⁴ ; Subj. N, r = 0.74, p = 1.1 × 10 ⁻ ²³; y: Subj. F, r = 0.60, p = 3.9 × 10 ⁻ ¹ ⁴ ; Subj. N, r = 0.73, p = 7.1 × 10 ⁻ ²³; Fig. 1h). Thus, the error structure in trained conditions predicted systematic biases in novel combinations.

The GP model also revealed a proximity effect in Start–Flow condition space. Its posterior uncertainty was correlated with the animals’ trial-to-trial saccadic variability (Subj. F, r = 0.23, p = 0.0095; Subj. N, r = 0.31, p = 3.9 × 10 ⁻ ⁴ ; Fig. 1i; Methods). Moreover, held-out conditions farther from the best-performing training condition with the same Flow direction, as measured by GP kernel distance, elicited larger errors (Subj. F, r = 0.31, p = 2.9 × 10 ⁻ ⁴ ; Subj. N, r = 0.30, p = 4.7 × 10 ⁻ ⁴ ; Fig. 1i; Methods). Together, these results indicate that generalization was more accurate and less variable near familiar conditions.

The proximity effect could arise if animals simply selected the response associated with the closest familiar condition: nearby Start–Flow combinations tend to have similar endpoints, making this strategy more accurate near trained conditions. To test this possibility, we compared each animal’s response distribution for each held-out condition with those for its five nearest trained conditions. For most held-out conditions (Subj. F, 80%; Subj. N, 76%), responses differed significantly from all five neighboring distributions (Fig. S1.4). Thus, selecting the response associated with a nearby trained condition could not explain most generalization behavior, supporting the interpretation that animals applied a learned computation to novel combinations.

### Characterizing the neural basis for compositional computation – Phase 2

The graded pattern of behavioral errors raised the question of how neural representations and dynamics support accurate composition, and whether distortions in this organization predict systematic errors. To address this question, we performed large-scale neural recordings in DMFC, which plays a central role in tracking variables through time using low-dimensional population dynamics ^18–20,24–29^.

Individual DMFC neurons exhibited diverse temporal responses modulated by Start and Flow in both Set 2a and Set 2b (Fig. 2a). Across the recorded populations (Subj. F: 1,005 units; Subj. N: 2,598 units), approximately 90% of units were selective for Start, Flow, or both, with more than 50% selective for both variables (Fig. 2b; Methods).

**Figure 2.**
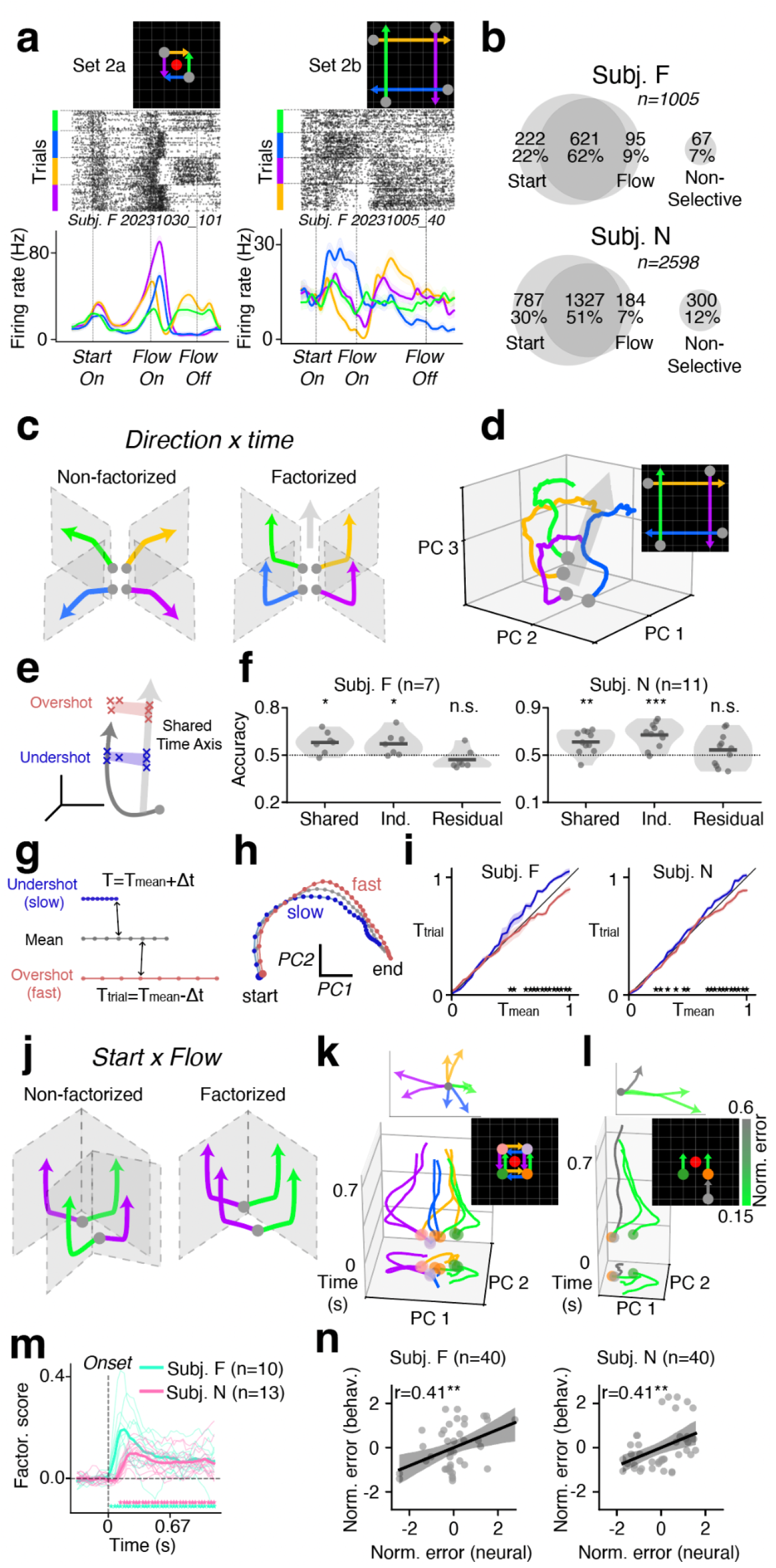
DMFC encodes task variables through behaviorally relevant factorized representations. **a**, Raster plot and peri-stimulus time histogram (PSTH) of example DMFC neurons for Set 2a (left) and Set 2b (right) conditions. Trials sorted by conditions (inset, color-coded). **b**, Population selectivity summary for Start, Flow, both, or neither. **c–f**, Factorization of Direction and Time. **c**, Schematics of non-factorized (left) and factorized (right) encoding. Non-factorized: Flow representations evolve differently for each direction. Factorized: they evolve along a shared temporal axis. **d**, Population trajectories (top 3 PCs) for four Flow directions (Set 2b, magnitude 6, Subj. F pseudo-population, *n* = 497). Conditions shown by inset. **e**, Schematic of overshot and undershot trials relative to the shared temporal axis. **f**, Accuracy of three decoding schemes for classifying over-versus undershoot trials (Methods). Shared, a single discriminant axis across all directions; Ind. (independent), separate axes per direction; Residual, independent decoding after projecting out the shared axis. Each dot is one session (*n*, number of sessions); bars show means. One-sided one-sample t-test against chance (0.5); *, p < 0.05; **, p < 0.01; ***, p < 0.001; n.s., not significant. **g–i**, KiNeT time-warping analysis. **g**, Schematic. For each time T_mean_ on the mean trajectory, the nearest state on the group-mean trial trajectory is found, yielding the matched trial time T_trial_ (Methods). **h**, Mean over-(red) and undershoot (blue) trajectories in the top 2 PCs (Set 2b, two subjects pseudo-population, *n* = 856). Magnitude 3 conditions averaged across directions. **i**, Matched trial time (T_trial_) versus mean-trajectory time (T_mean_) for over- and undershoot trajectories. Shading, s.e.m. across sessions. **, significant over–under separation (one-sided permutation test per time bin, FDR-corrected). **j–n**, Factorization of Start and Flow. **j**, Schematics of non-factorized (left) and factorized (right) encoding. Non-factorized: identical Flows drive different dynamics depending on Start. Factorized: identical Flows induce the same displacements regardless of Start. **k**, Population trajectories (top 2 PCs across time) for conditions varying in Start location and Flow direction (Set 2a, Subj. F pseudo-population, n=508). Trajectories color-coded by conditions (side inset). Top inset, the first 100 ms trajectories after subtracting the Start state. **l**, Same as k, including an upward Flow condition in which Subj. F performed poorly (color-coded by behavioral error). **m**, Additive factorization score (ΔR²) over time (Methods). Thin lines, individual sessions; thick lines, session means (*n*, number of sessions). *, ΔR² significantly greater than zero (one-sided one-sample t-test across sessions, FDR-corrected at α = 0.05). **n**, Neural deviation from factorized geometry versus normalized behavioral error (Methods). Each dot is one condition-to-template pair. Both axes z-scored. *n*, number of pairs. **, p < 0.01 for Pearson correlation.

To examine how these variables were organized across the population, we analyzed the Set 2a dataset that included each of four Start locations paired with each of the four Flow directions at magnitude 2. The Start and Flow subspaces were largely distinct (Fig. S2.1a). Both variables were decodable well above chance during their respective task epochs, although Start decoding declined modestly after Flow onset (Fig. S2.1b). Cross-temporal analysis further showed that both subspaces rotated over time, indicating that the population patterns encoding each variable changed as the trial progressed (Fig. S2.1c), consistent with prior work in frontal cortical areas ^18,20,30–33^. These results established that DMFC maintained information about both inputs through evolving population activity, providing a basis for examining how that activity supported their composition.

The task requires computing **s** + **d**t across combinations of Start, Flow direction, and Flow duration. We examined the neural organization of this computation in two steps. First, we used Set 2b, which varied duration within fixed Start–direction pairings, to ask how neural dynamics integrate Flow over time. Second, we used Set 2a, which varied Start and Flow direction at a fixed duration, to examine how the integrated displacement is combined with Start.

### Neural dynamics underlying temporal integration

Computing the displacement (**d**t) requires combining Flow direction with elapsed time. We considered two hypotheses for how neural dynamics implement this computation. In the first, separate integrators accumulate displacement for each of the Flow directions, producing four direction-dependent neural trajectories (Fig. 2c, left). In the second, direction and elapsed time are factorized: Flow onset shifts the neural state in a direction-dependent manner, after which activity evolves along a shared temporal dimension (Fig. 2c, right). This factorized scheme allows the same timing dynamics to be used across directions, with direction and time combined downstream to compute displacement.

To evaluate these hypotheses, we first visualized Set 2b population activity in the top three principal components, focusing on displacement magnitude 6 to reveal the trajectories over an extended Flow period (Figs. 2d and S2.2). The geometry was consistent with the factorized scheme: trajectories for different Flow directions were strikingly similar, appearing as shifted versions of one another for much of this period. This similarity extended even to opposite Flow directions (trajectory displacements from 300 ms to Flow offset with mean pairwise cosine similarity of 0.92; chance 95% CI: −0.31 to 0.57), suggesting that a common temporal component governed the evolution of activity across directions (Fig. 2d).

The shared timing axis makes a concrete prediction: if it tracks the animal’s estimate of elapsed time, trials that advance farther along it should correspond to overshoots, whereas trials that lag behind should correspond to undershoots (Fig. 2e). We tested this prediction with three complementary decoding analyses (Fig. 2f; Methods). First, we asked whether a decoder for the shared temporal dimension could predict whether the animal overshot or undershot the target; it could (one-sided one-sample t-tests against chance; F, n = 7 sessions: p = 0.010, d = 1.18; N, n = 11 sessions: p = 0.0014, d = 1.19). Second, we asked whether direction-specific decoders could do the same when applied separately to each Flow direction; they also could (F: p = 0.022, d = 0.96; N: p = 1.4 × 10⁻ ⁴, d = 1.65). However, once the shared temporal dimension was removed, the direction-specific decoders no longer performed significantly above chance (F: p = 0.87, d = −0.47; N: p = 0.16, d = 0.31). Together, these results support a shared timing dimension that captures magnitude errors across Flow directions, consistent with factorized representations of elapsed time and direction.

Previous work suggests that variability in perceived elapsed time reflects variability in the speed of neural trajectories ^19,20,34^. As a complementary test of the behavioral relevance of the shared timing dimension, we therefore asked whether trajectory speed predicted magnitude errors. We applied KiNeT ^18,20,25^, a time-warping analysis that compares the progression of neural trajectories through state space (Fig. 2g; Methods). We grouped trials into overshoots and undershoots and compared their group-mean trajectories with the mean trajectory. Overshoot trajectories advanced more rapidly than the mean trajectory, whereas undershoot trajectories advanced more slowly (Fig. 2h). Accordingly, the time required to reach a corresponding state on the mean trajectory fell below the identity line for overshoots and above it for undershoots (Fig. 2i; one-sided session-stratified permutation tests per time bin (1,000 permutations), FDR-corrected; F: 14/31 bins p < 0.05, n = 7 sessions; N: 17/31 bins p < 0.05, n = 11 sessions). These results link faster and slower neural dynamics to overestimation and underestimation of elapsed time, respectively. Together with the decoding analyses, they support a shared temporal dimension that represents elapsed time independently of Flow direction and whose variability predicts displacement magnitude errors.

### Combining integrated displacement with Start location

Next, we asked how the integrated displacement (**d**t) is combined with Start (**s**). Variation in performance across Set 2a allowed us to test whether accurate composition was associated with a factorized representation of these quantities. We hypothesized that accurate responses would be supported by neural trajectories in which the same Flow induces similar changes in activity across Start locations (Fig. 2j, right), whereas systematic errors would accompany Start-dependent distortions of those trajectories (Fig. 2j, left).

We first visualized the population activity as a function of time in the top two principal components for Set 2a, focusing on the four Start locations near fixation (Fig. 2k, S2.2). The trajectories formed four self-similar groups, each corresponding to one displacement, consistent with a factorized representation of Flow across Start locations. Subtracting the initial state associated with each Start and aligning trajectories to a common origin made this organization particularly evident early after Flow onset (Fig. 2k, inset). Notably, this parallel structure was disrupted in conditions with poorer performance (Fig. 2l).

A striking feature of this organization was that opposite Flow directions induced nearly opposite perturbations in neural state space: left opposed right, and up opposed down (Fig. 2k, inset). To quantify this geometry, we measured the cosine similarity between the perturbations for each pair of opposite directions over time. Mean cosine similarity (average of the left–right and up–down pairs) was strongly negative in both animals and more negative than expected for random mean-centered vectors (Subj. F: mean = −0.88, null 95% CI = [−0.71, 0.17]; Subj. N: mean = −0.74, null 95% CI = [−0.70, 0.16]; Fig. S2.1d). In conditions with poorer performance, this Cartesian structure was disrupted early after Flow onset, suggesting that its integrity was important for the ensuing integration (Fig. 2l, inset).

We quantified this structure by comparing the variance explained by an additive model that expressed population activity as separate contributions from Start and Flow with a shuffled baseline (shuffling Flow labels within each Start; Methods). This factorization score (ΔR²) significantly exceeded chance in both animals (Fig. 2m; Subj. F: ΔR² = 0.10 ± 0.02, n = 10 sessions, p = 1.6 × 10 ⁻ ⁴ ; Subj. N: ΔR² = 0.06 ± 0.01, n = 13 sessions, p = 7.0 × 10 ⁻ ⁶ ; mean ΔR² after Flow onset, mean ± s.e.m. across sessions, one-sided one-sample t-test). This structure emerged shortly after Flow onset and persisted through the delay period.

We then quantified the relationship between departures from this geometry and behavioral error by measuring each trajectory’s deviation from the geometry of the well-performing same-direction conditions (Methods). In both animals, larger neural deviations predicted larger behavioral errors (Fig. 2n; Subj. F: r = 0.41, p = 0.0089; Subj. N: r = 0.41, p = 0.0080), linking the integrity of the factorized organization to the accuracy of combining **d**t with **s**.

Together, these analyses show that DMFC activity evolves along a shared temporal dimension across Flow directions while maintaining a factorized representation of Start and Flow. Departures from this organization predict behavioral errors. We next asked whether the same organization supports composition in novel Start–Flow combinations.

### Structured neural dynamics supporting compositional generalization

Phase 3 challenged animals with novel Start–Flow combinations presented without visual guidance from the first trial. These conditions therefore allowed us to ask whether the neural organization established during training extended to new combinations of familiar inputs. We analyzed recordings from conditions that elicited either consistently accurate responses or systematic biases to identify the neural signatures of successful and unsuccessful generalization (Fig. S1.3).

Individual DMFC neurons again showed diverse task-dependent responses (Fig. 3a). As in Phase 2, many units were selective for Start, Flow, or both, with a substantial fraction selective for both variables (Fig. 3b; Methods). Thus, DMFC continued to encode both task inputs when animals encountered novel combinations.

**Figure 3.**
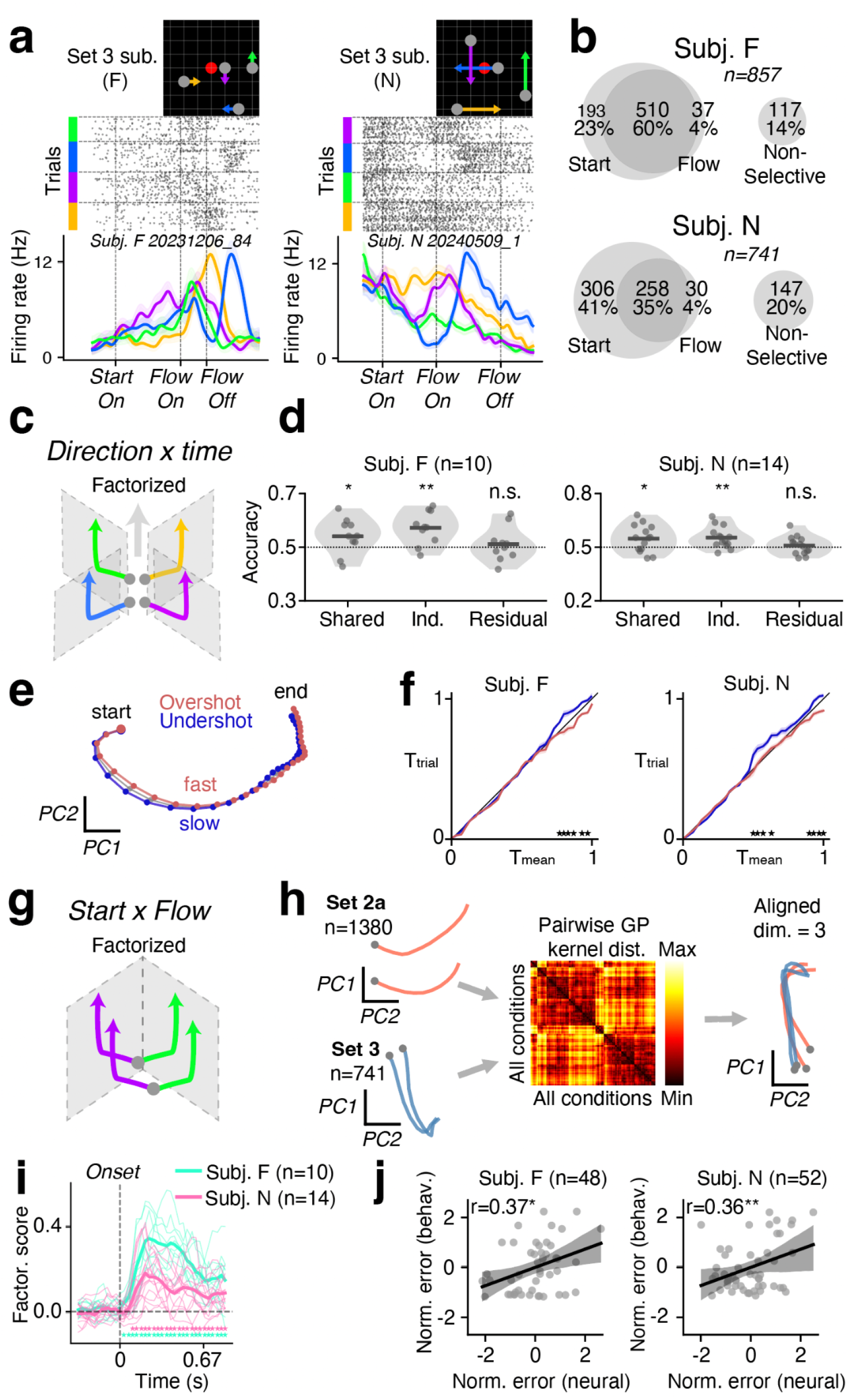
Factorized neural representations extend to novel Start × Flow combinations. **a**, Raster plot and PSTH of example DMFC neurons for Set 3 subset conditions in Subj. F (left) and Subj. N (right). Trials sorted by conditions (inset, color-coded). **b**, Population selectivity summary for Start, Flow, both, or neither for Set 3 sessions. **c–f**, Factorization of Direction and Time. **c**, Schematic of the factorized shared time encoding axis. **d**, Same three decoding schemes as Fig. 2f, applied to Set 3 (Methods). Each dot is one session (*n*, number of sessions); bars show means. One-sided one-sample t-test against chance (0.5); *, p < 0.05; **, p < 0.01; n.s., not significant. **e**, Mean over-(red) and undershoot (blue) trajectories in the top 2 PCs (Set 3, two subjects pseudo-population, *n* = 1,169 neurons). Magnitude 3 conditions averaged across directions. **f**, Matched trial time (T_trial_) versus mean-trajectory time (T_mean_) for over- and undershoot trajectories. Same KiNeT analysis as Fig. 2i. **g–j**, Factorization of Start and Flow in Set 3. **g**, Schematic of the factorized Start × Flow encoding hypothesis: identical Flows induce the same displacements regardless of Start. **h**, Kernel-aligned projection. Example trajectories (Subj. N) from Set 2a and Set 3 in their own PCA spaces and after projection into the shared subspace. Set 2a and Set 3 pseudo-populations are linearly projected into a shared low-dimensional subspace, optimized so that pairwise neural trajectory distances match GP kernel distances derived from Set 1 and Set 2 behavior (Methods). **i**, Additive factorization score (ΔR²) over time, same analysis as Fig. 2m applied to Set 3 (Methods). **j**, Neural deviation of Set 3 condition from Set 2a template condition (in the kernel-aligned subspace) versus behavioral error of the Set 3 condition (Methods). Each dot is one ‘Set 3 condition–Set 2a template’ pair. Both axes z-scored. *n*, number of pairs. *, p < 0.05; **, p < 0.01 for Pearson correlation.

We next asked whether the shared timing dimension continued to predict magnitude errors in Set 3 (Fig. 3c). The same decoding analyses used in Phase 2 yielded a similar pattern (Fig. 3d). A decoder for the shared temporal dimension distinguished overshoot from undershoot trials (one-sided one-sample t-tests against chance; F, n = 10 sessions: p = 0.043, d = 0.61; N, n = 14 sessions: p = 0.017, d = 0.63). Direction-specific decoders also captured this variability (F: p = 0.0026, d = 1.16; N: p = 0.0024, d = 0.90), but no longer performed significantly above chance after the shared temporal dimension was removed (F: p = 0.30, d = 0.18; N: p = 0.26, d = 0.17). KiNeT analysis likewise showed that overshoot trajectories progressed faster than the mean trajectory, whereas undershoot trajectories progressed more slowly (Fig. 3e,f; one-sided session-stratified permutation tests per time bin (1,000 permutations), FDR-corrected; F: 6/31 bins p < 0.05, n = 10; N: 8/31 bins p < 0.05, n = 14). Thus, a shared, behaviorally relevant timing dimension was preserved across Flow directions even in novel Start–Flow combinations.

We next asked whether Start and Flow remained factorized in the generalization set (Fig. 3g). Because Set 3 was not designed to systematically cross Start and Flow, we assessed its neural trajectories against the structure identified in Set 2a. This comparison posed a challenge: the two sets contained different conditions and were recorded from different neurons. To overcome this, we developed a method that uses behavioral similarity to align neural datasets without requiring shared neurons or matched task conditions. Specifically, the GP fitted to Set 1 and Set 2 behavior defined a geometry of task conditions that served as a common reference for both neural datasets. We then learned linear projections of the two pseudo-populations into a shared latent space, matching distances between neural trajectories to those prescribed by the GP kernel (Methods). This approach allowed us to compare trained and novel neural trajectories within a common geometry anchored in behavior.

We first visualized example trajectories from Set 2a and Set 3 in their respective PCA spaces (Fig. 3h). After alignment, pairwise trajectory distances in the shared latent space closely matched those prescribed by the GP kernel (Subj. F: r = 0.80; Subj. N: r = 0.95; Fig. S3.1), showing that the linear projections could bring the two datasets into the common geometry defined by behavior. Within Set 3, the factorization score used in Phase 2 again revealed additive Start–Flow structure (Fig. 3i; Subj. F: ΔR² = 0.11 ± 0.01, n = 10 sessions, p = 4.0 × 10 ⁻ ⁷ ; Subj. N: ΔR² = 0.05 ± 0.01, n = 14 sessions, p = 3.1 × 10 ⁻ ⁵ ; same statistics as in Fig. 2m), indicating that this organization extended to novel combinations.

We then used the aligned space to ask whether departures from the trained dynamics predicted generalization errors. As in the Phase 2 analysis, we compared each Set 3 trajectory with a well-performed Set 2a template sharing the same Flow direction. Greater deviation from the trained template was associated with larger behavioral errors (Subj. F: r = 0.37, p = 0.010; Subj. N: r = 0.36, p = 0.0079; Fig. 3j). This relationship was robust across choices of latent dimensionality (Fig. S3.2).

Could the correlation between neural deviation and behavioral error follow simply from aligning neural activity with a kernel that already predicts behavior? To test this possibility, we shuffled Set 3 trajectories among conditions with the same Flow magnitude and realigned the shuffled neural data using the same GP kernel. If the correlation arose from the kernel-guided alignment alone, it should persist after shuffling and realignment. The observed correlations were stronger than those obtained after shuffling (1,000 permutations; Subj. F: p = 0.044; Subj. N: p = 0.030; Fig. S3.3), supporting a link between neural dynamics and behavioral errors beyond that imposed by the alignment.

### Architectural requirements for compositional computation and its failures

Our results identified three properties of vector composition. First, activity evolved along a shared temporal dimension across Flow directions, and trial-to-trial variation along this dimension predicted magnitude errors (shared-axis; Figs. 2d–i and 3c–f). Second, neural deviations from trained template trajectories covaried with behavioral errors across conditions: larger departures from the trained structure accompanied less accurate composition (error-coupling; Figs. 2n and 3j). Third, behavioral errors increased with distance from trained conditions in task space (proximity; Fig. 1g–i).

What circuit organization could give rise to these three properties, and what inductive biases would it require? To address these questions, we used recurrent neural network (RNN) models, which provide a framework for linking circuit connectivity to neural dynamics and computation ^3,17–20,35–45^. We developed RNNs with different internal organizations and asked which architectural features were sufficient to reproduce all three experimental signatures (Fig. 4). All models shared the same input–output format: four direction-specific inputs conveyed Flow for its duration, a 49-dimensional retinotopic input encoded Start throughout the trial, and a two-dimensional output specified the saccade target in Cartesian coordinates. Models were trained on the same 108 conditions used to train the animals (Sets 1 and 2) and evaluated on the remaining holdout conditions across multiple random initializations (Methods).

**Figure 4.**
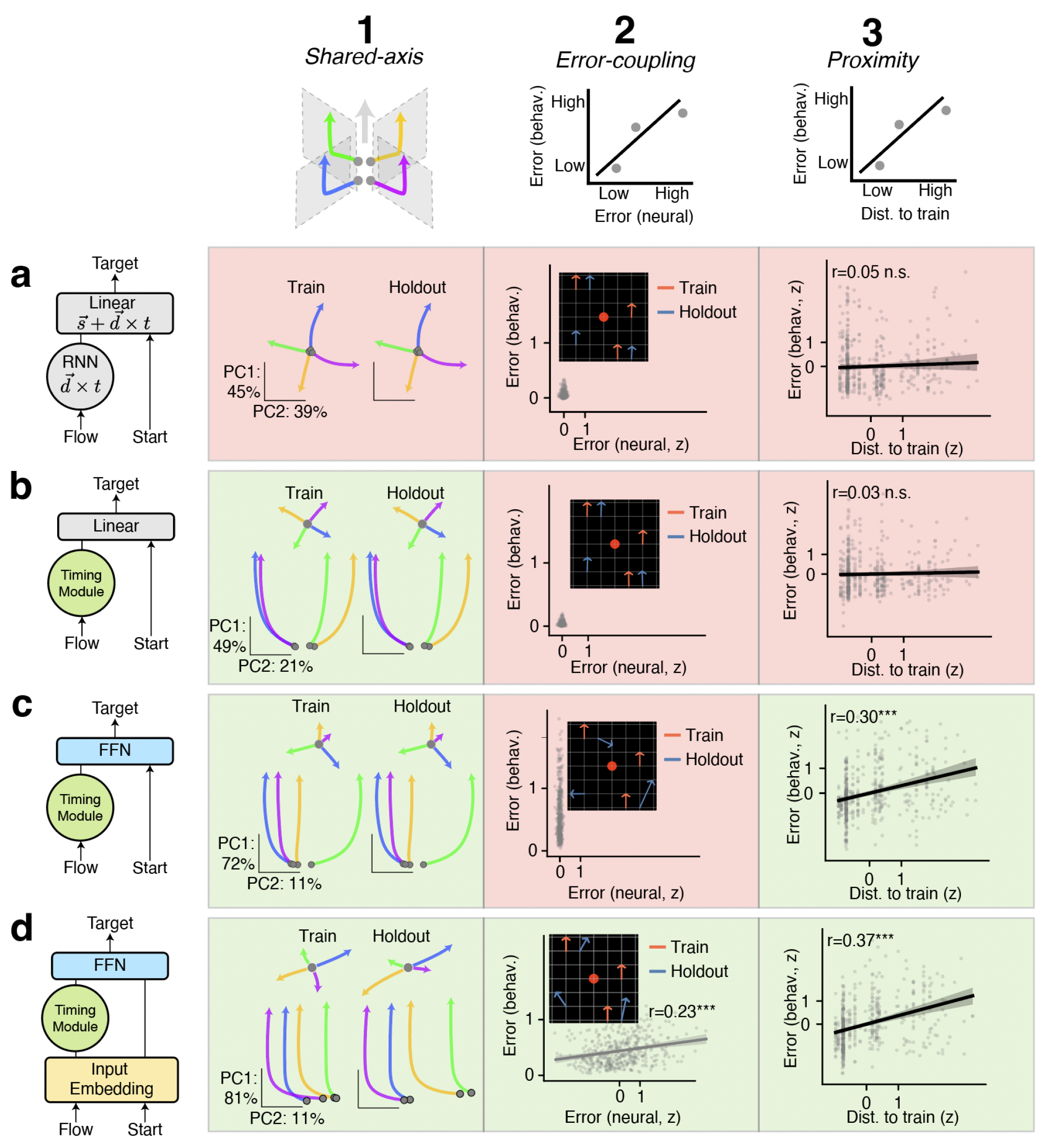
Architectural constraints inferred from neural network models. Four recurrent network architectures (a–d) evaluated against three features of the monkey’s vector composition (columns 1–3). Top row, schematic of the three features. Column 1, Shared axis for time-integration: PCA of integrator hidden states for four flow directions of magnitude = 6. Inset, the mean-removed trajectories subtracting Start state. Training and holdout conditions are plotted separately. Column 2, neural–behavioral correlation: neural trajectory deviation versus behavioral error for holdout conditions. Inset, example saccades of training and holdout conditions. r, Pearson correlation. Column 3, proximity effect: distance to nearest training condition versus behavioral error. r, Pearson correlation. Green background indicates the model replicates the monkey feature; red indicates failure. **a**, Flow drives an unconstrained RNN; hidden state is combined with Start through a linear output layer. **b**, Flow drives a Timing Module (pre-trained, frozen recurrent weights); hidden state is concatenated with Start and passed through a linear layer. **c**, Same as b, but the hidden state is concatenated with Start and passed through a nonlinear feedforward network (FFN). **d**, Same as c, but with a nonlinear input embedding that jointly transforms Flow and Start before they enter the two-stream architecture.

### A task-aligned architecture

We began with an architecture designed to directly implement the function **s** + **d**t (Fig. 4a). Flow entered a recurrent network that integrated the directional input over time to compute displacement dt. A linear readout then added this displacement to Start s to produce the target location.

The model generalized almost perfectly to held-out conditions, but its internal dynamics differed from those observed in DMFC. During integration, leftward and rightward Flow drove activity along approximately opposite trajectories, as did upward and downward Flow, reflecting the Cartesian geometry of the displacement vectors (Fig. 4a, left). This resembles a mental simulation that tracks displacement as it unfolds. In DMFC, by contrast, opposite Flow directions produced opposing perturbations early after Flow onset, but the ensuing trajectories evolved along a shared temporal dimension. Thus, the distinction concerned how activity evolved during integration: the model tracked direction-dependent displacement, whereas DMFC represented elapsed time through dynamics shared across directions.

The model also failed to reproduce the animals’ two characteristic error signatures. With Flow and Start entering through separate pathways and combined through a linear readout, neural deviations and behavioral errors remained near zero across held-out conditions, leaving no graded neural–behavioral error relationship (Fig. 4a, middle). Accuracy was also essentially independent of distance from the training set, so no proximity effect emerged (Fig. 4a, right). Thus, directly implementing the task’s computation supported nearly perfect generalization but reproduced neither the shared temporal dynamics nor the structured errors observed in the animals.

### A dedicated timing module

The task-aligned model suggested that reproducing the shared temporal dynamics observed in DMFC would require an additional inductive bias. We reasoned that the model might have adopted separate integration trajectories for the four Flow directions because full-rank recurrent connectivity afforded sufficient dimensionality. Restricting the rank might therefore force integration onto a shared axis. We tested this possibility by training the same task-aligned architecture with different ranks of recurrent connectivity. However, even rank-1 networks used nonlinearities to generate direction-specific integration trajectories (Fig. S4.1).

We therefore considered whether DMFC might rely on a dedicated timing mechanism whose dynamics are shared across directions. We pre-trained an RNN to integrate a constant input into a monotonic estimate of elapsed time, then froze its recurrent weights and incorporated it into the task-aligned architecture (Fig. 4b). Flow drove this Timing Module, and its recurrent state was combined with Start through the same linear readout.

The Timing Module successfully reproduced the shared temporal geometry observed in DMFC: despite differences in Flow direction, neural activity evolved along a common temporal axis (Fig. 4b, left). Thus, pretraining and preserving a dedicated timing mechanism provided an inductive bias sufficient to generate this organization.

However, the model did not use this shared temporal signal to estimate displacement. Perturbing activity along the common timing axis had little effect on the predicted displacement magnitude; instead, the model relied on residual direction-specific components orthogonal to the shared axis to compute magnitude (Methods; Fig. S4.2). The model therefore reproduced the shared temporal geometry, but not its behavioral relevance: in DMFC, variation along the shared timing dimension predicted overshooting and undershooting.

The model also failed to reproduce the two remaining signatures. Neural deviations and behavioral errors remained near zero across held-out conditions, leaving no graded neural–behavioral error relationship (Fig. 4b, middle). Accuracy was independent of distance from the training set, so no proximity effect emerged (Fig. 4b, right). A dedicated timing mechanism therefore reproduced one aspect of the neural organization, but additional architectural changes were needed to capture how that organization relates to behavior.

### A nonlinear readout

The near-perfect generalization of Models a and b suggested that their task-aligned linear readout prevented the distance-dependent errors observed in the animals. We therefore replaced it with a nonlinear feedforward network while retaining the pretrained Timing Module and separate Start and Flow pathways (Fig. 4c). The feedforward network received the Timing Module’s recurrent state together with Start and produced the target location.

We first asked whether replacing the linear readout with a nonlinear network altered the shared temporal geometry. It did not: activity continued to evolve along a common timing axis across Flow directions (Fig. 4c, left). We then asked whether the nonlinear readout made this shared timing signal behaviorally relevant. It did: unlike Model b, perturbing activity along the common axis changed the predicted displacement magnitude (Methods; Fig. S4.2). Thus, Model c preserved the shared temporal dynamics while allowing the readout to use them to estimate displacement.

We next asked whether Model c reproduced the proximity effect. In Models a and b, the linear readout matched the additive structure of the task and supported nearly perfect generalization. In Model c, the readout instead learned a general nonlinear mapping from the training examples, without an architectural constraint enforcing addition. The model performed accurately on trained conditions but made errors on held-out combinations. These errors increased with distance from the training set, reproducing the proximity effect observed in the animals (Fig. 4c, right).

However, Model c did not reproduce the coupling between neural deviations and behavioral errors. Because the Start and Flow inputs remained factorized, the same Flow drove identical recurrent dynamics across Start locations, including in novel combinations. Neural deviation therefore remained zero even when behavioral errors were large (Fig. 4c, middle). Thus, the model captured the behavioral relevance of the shared timing signal and the proximity effect, but its factorized inputs prevented the trajectory distortions that accompanied errors in the animals.

### Entangled inputs

Model c produced behavioral errors without distorting the recurrent trajectories, suggesting that reproducing their coupling required a source of distortion before the readout. In DMFC, Flow onset produced direction-dependent perturbations that respected the Cartesian geometry of the task (Fig. 2k, inset; Fig. S2.1d), and departures from the factorized trajectory structure predicted behavioral errors (Fig. 2n). Errors introduced solely by a downstream readout could not explain these deviations in the recorded dynamics.

We therefore asked whether entangling Start and Flow before they reached the Timing Module would reproduce this relationship. Such mixing could arise as spatial and motion signals in visual cortical areas undergo transformations before reaching DMFC. We added a nonlinear embedding layer that jointly transformed Start and Flow, leaving the remaining architecture unchanged from Model c (Model d; Fig. 4d).

This model reproduced all three experimental signatures. The frozen Timing Module preserved the shared temporal axis across Flow directions (Fig. 4d, left). Because the inputs were no longer fully factorized, the same Flow drove different trajectories depending on Start, and novel combinations produced neural deviations that covaried with behavioral errors (r = 0.23, p = 3.1 × 10 ⁻ ⁷) (Fig. 4d, middle). The model also retained the proximity effect, with larger errors farther from the training set (r = 0.37, p = 7.4 × 10 ⁻ ¹ ⁷) (Fig. 4d, right).

### An alternative locus for the entanglement

Model d introduced entanglement before the Timing Module, but Start and Flow could also become mixed within the recurrent dynamics. To test this possibility, we constructed a model in which both inputs entered the Timing Module directly, leaving the remaining architecture unchanged (Fig. S4.3). This model also reproduced all three experimental signatures, indicating that our findings are consistent with entanglement arising either before or within DMFC.

Together, these models linked the three experimental signatures to specific architectural features. A dedicated timing mechanism reproduced the shared temporal geometry, and a nonlinear readout allowed that timing signal to guide behavior while introducing distance-dependent errors. Reproducing the coupling between neural deviations and behavioral errors additionally required entanglement of Start and Flow before or within the Timing Module. These findings suggest that compositional generalization depends on alignment between input representations, the dynamics that transform them, and the operations that map neural states onto behavior.

### Testing shared principles of compositional generalization in humans

The monkeys’ graded failures are puzzling given how straightforward the task feels to us: once we know a displacement, applying it from a new starting point seems trivial. What accounts for this difference? The mapping from optic flow around fixation to displacement from a separate Start location may be intuitive to humans but unfamiliar to monkeys. Our models offer a possible explanation. The task-aligned architecture generalized almost perfectly (Fig. 4a), whereas architectures with entangled inputs or an output mapping that departed from the task’s additive structure exhibited graded failures (Fig. 4c,d). This comparison suggests that our apparent ease may depend on an established alignment between sensory inputs and the operations required to produce a response. Humans might therefore exhibit similar failures when that alignment is unfamiliar and must be learned.

To test this hypothesis, we designed three experiments that varied the mapping between the Flow stimulus and the required transformation (Fig. 5a,b). Experiments 1 and 3 used congruent mappings. In Experiment 1, Cartesian Flow specified Cartesian displacement, as in the monkey task. In Experiment 3, both the stimulus and the required displacement were polar: radial Flow specified radial motion, and angular Flow specified angular motion. Experiment 2 instead paired Cartesian Flow with polar displacement, requiring subjects to learn a nonintuitive association between familiar stimuli and transformations. We predicted that subjects would generalize accurately in Experiments 1 and 3 but show graded failures in Experiment 2, where the sensory cues were not aligned with the required computation.

**Figure 5.**
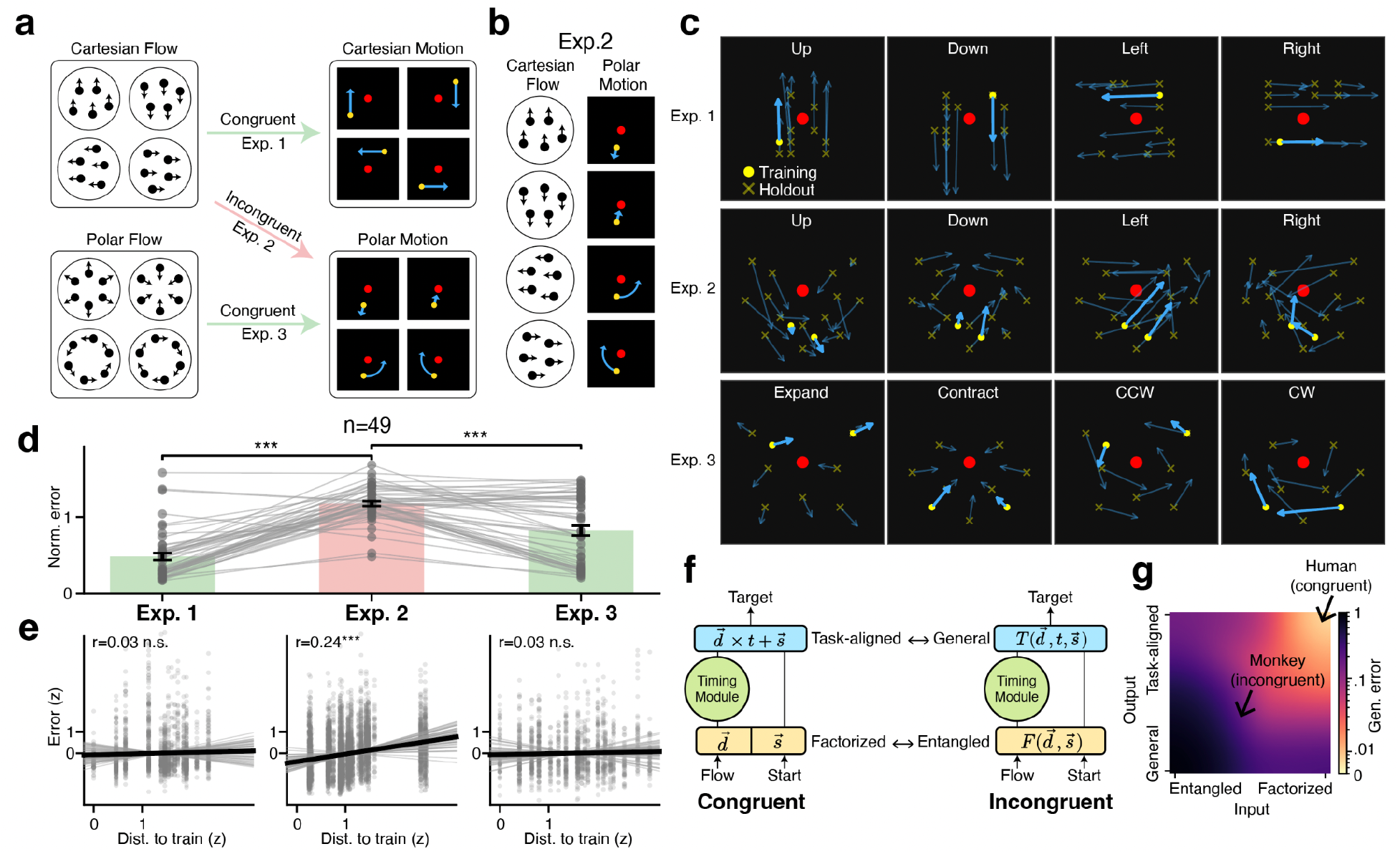
Proximity effect in human compositional generalization. **a**, Experimental design. Subjects performed three experiments in a single session. Experiment 1: Cartesian flow mapped to Cartesian motion (congruent). Experiment 2: Cartesian flow mapped to polar motion (incongruent). Experiment 3: polar flow mapped to polar motion (congruent). **b**, Incongruent mapping in Experiment 2. Each Cartesian flow direction (left column) maps to a polar transformation (right column): up → expand, down → contract, left → CCW, right → CW. **c**, An example subject’s responses across all three experiments. Each panel shows one flow condition; arrows point from start to mean click location. Yellow circles, training positions; yellow crosses, holdout positions. Red circle, screen center. Training responses are opaque; holdout responses are translucent. **d**, Normalized error on holdout conditions across experiments (*n* = 49 subjects). Bars, population means and s.e.m.; each dot is one subject; gray lines connect individual subjects. ***, p < 0.001 (paired t-test). **e**, Proximity analysis. Distance to nearest training condition (z-scored) versus holdout error (z-scored) for each experiment. Gray dots, individual trials; thin lines, per-subject regressions; thick line, pooled regression. r, Pearson correlation; ***, p < 0.001; n.s., not significant. **f**, Model architectures for congruent (left) and incongruent (right) solutions. Congruent: factorized Flow feeds a Timing Module; output composed through a task-aligned feedforward network (FFN, for multiplication) and linear layer (for addition with Start). Incongruent: entangled Start and Flow input embedding feeds a Timing Module; output mapped through a general-purpose FFN. **g**, Generalization error as a function of input factorization (x-axis) and output task-alignment (y-axis), obtained by continuously interpolating between model architectures (Methods). Arrows indicate the regimes corresponding to human congruent performance (top right, near-zero error) and monkey/human incongruent performance (intermediate region, graded error).

Subjects completed all three experiments online within a single session. In each experiment, they trained on a small subset of Start locations with visual guidance and were then tested on held-out locations without guidance (Methods). In Experiment 2, training was concentrated near the bottom of the display, where the required polar displacement was strongly incongruent with Cartesian Flow. Held-out locations sampled increasing distances from the training region, allowing us to test whether generalization showed the distance-dependent errors observed in monkeys.

The behavioral contrast was strong and evident within individual subjects. In Experiment 1, subjects accurately applied the Cartesian displacements specified by Flow across trained and held-out Start locations (Fig. 5c, top). They also generalized the radial and angular transformations in Experiment 3 (Fig. 5c, bottom). In Experiment 2, however, responses showed systematic errors, particularly away from the trained conditions (Fig. 5c, middle). Across subjects, normalized error was significantly greater in Experiment 2 than in either congruent experiment (paired t-tests; n = 49; Exp. 1 versus Exp. 2, t(48) = −13.6, p = 4.2 × 10⁻¹ ⁸ ; Exp. 2 versus Exp. 3, t(48) = 6.6, p = 2.9 × 10⁻ ⁸ ; Fig. 5d).

Remarkably, errors in Experiment 2 showed the same proximity effect observed in monkeys. Behavioral error increased systematically with distance from trained conditions (mean per-subject r = 0.24 ± 0.02 s.e.m., one-sample t-test, t(48) = 11.9, p = 5.6 × 10 ⁻ ¹ ⁶ ; Fig. 5e; Methods). Neither congruent experiment showed a significant proximity effect (Experiment 1: r = 0.03 ± 0.02, p = 0.085; Experiment 3: r = 0.03 ± 0.02, p = 0.18; Fig. 5e). This pattern could arise from a memorization strategy in which subjects responded to held-out conditions using responses to similar training conditions. However, further analysis indicated that memorization alone could not account for the proximity effect (Fig. S5.2; Methods).

The variability of human subjects across congruent and incongruent conditions and our earlier modeling results suggest that task alignment may lie on a continuum. Input representations can be factorized to different degrees, and output operations can vary in how closely they match the required computation. We therefore asked how these two dimensions of alignment jointly shape compositional generalization.

To examine this continuum, we constructed an architecture in which *input factorization* and output *task-alignment* could be varied independently while retaining the same Timing Module (Fig. 5f; Methods). At the input, we varied the degree of entanglement between Start and Flow. At the output, we varied how displacement and Start were combined, spanning a general nonlinear mapping and a linear operation matched to the task’s additive structure. At the fully aligned endpoint, a nonlinear module converted direction and elapsed time into displacement, which was then added linearly to Start. This organization allowed the shared timing signal to guide behavior while supporting accurate generalization (Figs. S4.2 and S5.3).

Generalization varied systematically across this continuum (Fig. 5g). Errors were largest when inputs were strongly entangled and the output implemented a general nonlinear mapping. As input representations became more factorized and the output operation more closely matched the task, errors declined, approaching zero when both were aligned. These simulations show how varying degrees of alignment at the input and output can produce a graded range of generalization, from task-aligned composition to distance-dependent failures observed in our experiments. These results suggest that alignment between input representations, dynamics and readout operations is critical for compositional generalization.

## Discussion

Research over the past decade has converged on the idea that cognitive flexibility relies on structured, reusable neural resources. Cognitive maps enable flexible navigation through physical space ^46,47^, non-spatial variables ^16,22,48,49^, and abstract spaces without sensory input ^14^. Structured representations support generalization across contexts ^9,10^, tracking progress through goals ^5^, and reuse of sensory and motor components across tasks ^1^. Slot-like codes likewise organize sequence memory ^13^, and invariant action representations enable recombination into novel action sequences ^6^. Reuse also extends to dynamical motifs ^3,37^ across motor ^50,51^, timing ^17,19^, and cognitive tasks ^18,20^. Together, these findings set the stage for elucidating how novel combinations of existing representations and dynamics support compositional generalization ^23,52^.

Our study begins to address this problem by examining how recurrent dynamics acting on novel combinations of inputs support correct computation in regions of state space not visited during training. We designed a vector composition task in which animals learned all individual input values but only a subset of their combinations, allowing us to test the learned operation on combinations withheld from training. Recordings in DMFC revealed shared temporal dynamics across Flow directions and a factorized organization of Start and Flow that supported accurate composition. Importantly, there were also conditions in which animals’ behavior systematically deviated from the desired composition. These failures allowed us to examine the nature of behavioral errors and the corresponding departures from the underlying neural organization. Comparing these patterns of success and failure with circuit models revealed that compositional generalization poses a three-way alignment problem: input representations, recurrent dynamics, and readout operations must work together to implement the required computation.

An intuitive solution would be to mentally simulate a point moving from Start in the direction specified by Flow, continuously tracking its position until Flow offset. Our unconstrained RNNs demonstrated that such a solution was viable, integrating displacement along direction-specific trajectories. DMFC exhibited a different organization: Start established the initial state, Flow direction shifted that state within a Cartesian organization, and activity then evolved along a shared temporal dimension. This arrangement preserved the components needed to compute the endpoint without requiring the temporal dynamics themselves to trace its changing spatial position. The endpoint could then be obtained by combining Start, direction, and elapsed time through an appropriate readout. Although this organization does not exclude an explicit endpoint representation elsewhere in the brain, it reveals the importance of the brain’s internal timing mechanisms as a building block of the underlying computational strategy.

A key finding was the presence of self-similar neural trajectories with different initial states established by Start and Flow. Notably, the speed of activity along these trajectories predicted magnitude errors. Together with prior work implicating such dynamics in a variety of timing tasks ^17–20,27–29,53,54^, these findings suggest that animals used a timing mechanism to estimate vector magnitude from Flow duration. Whether this mechanism preceded task learning or emerged through training remains an open question.

Representing elapsed time separately from direction requires a readout that combines time, direction, and Start according to the task’s multiplicative and additive operations. In our models, a linear readout could not use the shared timing signal for this purpose and instead relied on direction-specific components of activity. A nonlinear readout made the shared signal useful for estimating displacement, but accurate generalization also depended on how that displacement was combined with Start. Thus, recruiting a timing mechanism for spatial composition requires a readout matched to both its representational format and the task’s operations.

Recruiting this timing mechanism also requires appropriately structured inputs. Start and Flow information must reach DMFC through sensory pathways. Start information is widely available across retinotopic representations, and Flow drives direction-selective neurons in motion-sensitive areas such as MT. Although sensory processing makes these signals separately available, intervening nonlinear transformations may entangle them before they reach DMFC. We found that accurate composition was associated with factorized Start and Flow representations. In these conditions, Flow-induced shifts also exhibited a Cartesian organization, with opposing shifts for opposite directions consistent with motion opponency in motion-sensitive areas. In contrast, departures from factorization and Cartesian organization accompanied systematic errors, consistent with the effects of entanglement in our models. Learning to compose may therefore depend partly on routing and transforming sensory information into a format compatible with the existing recurrent computation.

The systematic biases reveal how this coordination extends beyond trained conditions. Behavioral biases varied smoothly across condition space, allowing us to use a Gaussian process fitted to errors on trained conditions to predict errors on novel combinations. Departures from the neural geometry supporting accurate composition likewise predicted behavioral error. Together, these findings suggest that the computation remained approximately valid near familiar conditions but became systematically distorted as inputs moved farther away. The models provide a possible mechanism for this pattern: varying input factorization and output alignment produced a continuum from nearly exact composition to persistent, distance-dependent biases. Generalization therefore depended on how far the coordination established through learning extended beyond the conditions that shaped it.

The human experiments suggest that these systematic failures are not unique to monkeys. Humans generalized accurately when sensory cues were congruent with the required transformations, but developed distance-dependent biases when familiar cues specified familiar operations through an unfamiliar association. The difficulty therefore could not be attributed simply to an inability to perform Cartesian or polar transformations. Instead, it arose when the relationship between the cue and the required computation had to be established anew. This result suggests that the apparent ease of familiar compositions may depend on alignment acquired through prior experience. The human data support the relevance of this computational constraint across species, although they do not establish that humans used the same neural implementation observed in DMFC.

Several questions remain about how the brain establishes this alignment. Our recordings linked changes in neural geometry to behavioral errors, and our models showed how those changes could arise. However, more than one circuit could explain the findings. Start and Flow could become entangled before reaching DMFC or within its recurrent circuitry. We also do not know where neural activity is converted into the endpoint used to guide behavior. Recording during learning could reveal whether generalization improves as inputs become more factorized, dynamics change, or the readout becomes better matched to the task. Perturbing neural activity would offer a more direct test: shifting activity along the shared timing dimension should change the estimated displacement magnitude across Flow directions, whereas shifting its direction-related components should change how that timing signal is translated into spatial displacement.

Compositional generalization thus depends on more than the availability of reusable representations and computations. Those resources must remain coordinated when unfamiliar inputs engage the circuit. By examining successful compositions together with systematic failures, our results connect the reliability of a learned operation to the alignment between its input representations, internally generated dynamics, and readout.

## Methods

### Subjects

Experimental procedures for animals conformed to the National Institutes of Health guidelines and were approved by the Committee on Animal Care at the Massachusetts Institute of Technology. We collected behavioral and neurophysiological data from two male rhesus macaque monkeys (Macaca mulatta, ID: F and N, age: 12 and 8 years old, weight: 13 and 12.5 kg).

### Experimental setup

Monkeys were head-restrained and seated comfortably in a dark, quiet room. Visual stimuli were presented on a 58-cm (23-inch) monitor (60 Hz refresh rate) positioned 60 cm from the animals’ eyes. Eye position was recorded using an infrared eye-tracking system (EyeLink 1000, SR Research) sampled at 1 kHz. Stimulus presentation and the registration of eye position were controlled using the MWorks software package (https://mworks.github.io/). A photodiode was used to precisely record the timing of visual stimulus events.

### Task structure

Monkeys were trained to perform a vector addition task. Visual stimuli were presented on a 7 × 7 square grid with coordinates [− 3, − 2, − 1, 0, 1, 2, 3] on X and Y axes (grid lines are visible). The grid spanned 24° × 24° of visual angle, with each grid cell measuring 4° × 4°. Grid intersections defined the discrete spatial locations used in the task. The distance between adjacent grid intersections was defined as 1 grid unit.

Each trial began when the animal fixated a red fixation point located at the center of the grid (coordinates [0, 0]). Upon successful fixation, a yellow point appeared at one of the grid intersections, indicating the Start location. The animal was required to maintain fixation for 0.83 s, after which a Flow cue was presented around the fixation point.

The Flow stimulus consisted of a coherent motion field with a circular aperture of radius 1 grid unit and a speed of 3 grid units per second. The duration of the Flow stimulus determined the displacement vector, defined as the product of motion direction, speed, and duration. Following Flow offset, a variable delay period (0.4–0.7 s) ensued during which the animal was required to maintain fixation.

The Go cue was signaled by the disappearance of the fixation point. The animal then had up to 4 s to make a response. A response was registered when the animal executed a saccade to a grid intersection and maintained gaze within a radius of 0.4 grid units for at least 0.4 s. Only the first registered response was counted.

The correct response (Target) was defined as the vector sum of the Start location and the Flow-integrated displacement vector. Juice reward was maximal when the response matched the Target location and decreased linearly to zero as the error increased, reaching zero when the error equaled half the length of the displacement vector. Trials were aborted if fixation was broken before the Go cue. Neither the Start nor the Target location was allowed to coincide with the fixation point.

During training, visible trials were interleaved with invisible trials. In visible trials, a blue dot moved from the Start location toward the Target location during the Flow period at a speed matched to the Flow stimulus. The animal could solve the task by tracking this moving cue. Following successful completion of a visible trial, the opacity of the blue dot for that condition was reduced by 5% on subsequent presentations; conversely, errors increased the opacity by 5%. Set 3 conditions were presented only as invisible trials.

### Gaussian Process model of behavior

The Gaussian Process model was fitted on trial-averaged Set 1 and Set 2 behaviors (invisible conditions only) and was used to predict holdout conditions in Set 3. Training sessions were the last five sessions of Set 2a and Set 2b where behavior had stabilized (Set 1 conditions were interleaved in those sessions); the Set 3 sessions were the first five sessions of Phase 3. For each condition, we computed the mean saccade across sessions *m* = (⟨*e_x_* ⟩, ⟨*e_y_* ⟩) and the trial-to-trial endpoint variability

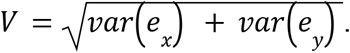

We modeled the saccade error field — the mean spatial deviations from Target — as a Gaussian process (GP) over the four-dimensional condition space *c* = (*x*_0_, *y*_0_, Δ*x*, Δ*y*): the Start position (*x*_0_, *y*_0_) and the Flow displacement (Δ*x*, Δ*y*). The dependent variable was the residual error *r* = *m* − *g*, where *g* is the ideal Target. We fit two independent GPs, one for each spatial component (*r_x_* and *r_y_*).

The covariance function was a Matérn-5/2 kernel:

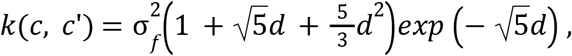

where 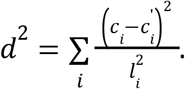.

The Matérn-5/2 kernel produces twice-differentiable sample functions, matching the expected smoothness of oculomotor error fields while avoiding the infinitely smooth assumption of the squared-exponential kernel. The ARD parameterization allows each condition-space dimension to contribute a different spatial scale to the covariance. The free parameters were the signal amplitude σ*_f_*, four dimension-specific lengthscales 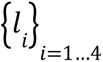 (one for each of *x*_0_, *y*_0_, Δ*x*, Δ*y*).

We minimized the standard GP negative log marginal likelihood with complexity regularization ^55^:

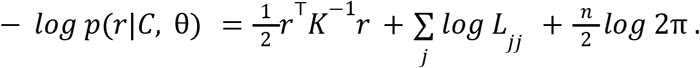

Term 1 (data fit): the Mahalanobis distance of the observed residuals *r* under the model covariance. The noisy covariance matrix *K* was constructed by adding observation noise to the kernel matrix: 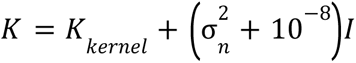. *K_kernel_* is the *n*×*n* matrix of kernel evaluations *k*(*c*, *c*’) between all training conditions, 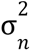 is a free parameter for the observation noise variance, *I* is the identity matrix.

Term 2 (complexity): the log-determinant of *K*, computed as the sum of log-diagonal entries of its Cholesky factor *L* (where *K* = *LL*^T^) — penalizes overly flexible models.

Term 3 (normalization): a constant depending only on the number of training conditions *n*.

Free parameters were optimized by minimizing this objective using L-BFGS-B (SciPy 1.12, https://scipy.org/) with five random restarts, retaining the solution with the lowest value.

Given the optimized hyperparameters, the posterior mean and variance at holdout conditions *C* were computed via standard GP equations:

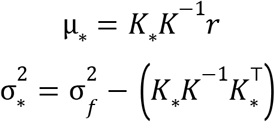

Where *K*_*_ is the cross-covariance between training and holdout samples. The combined posterior standard deviation was defined as 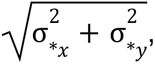, pooling the independent predictions for the *x* and *y* components.

The GP posterior mean 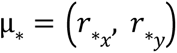 at each generalization condition was compared against the observed mean residual error 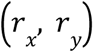. We report Pearson correlations of the *x* and *y* components separately in Fig. 1h.

The combined GP posterior standard deviation was compared against the trial-to-trial saccade variability *V* of each Set 3 condition in Fig. 1i left.

We computed the kernel distance between two conditions as:

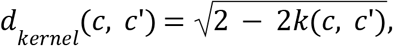

converting each pair of 4-dimensional condition vectors *c* = (*x*_0_, *y*_0_, Δ*x*, Δ*y*) into a scalar, measuring their distance in the GP space, where *k* is the Matérn-5/2 kernel evaluated with unit amplitude and the fitted GP lengthscales, so that *k*(*c*, *c*) = 1 (identical conditions yield distance of 0 and dissimilar conditions yield positive values).

In Fig. 1i right, for each holdout condition in Set 3, we first identified the training condition of the same Flow direction that achieved the lowest normalized error (indicating ideal learning). We then computed the kernel distance between the holdout condition and the ideal training condition, and compared against the actual error of the holdout condition.

### Neurophysiology

We recorded neural activity from the dorsomedial frontal cortex (DMFC) using a Neuropixels 1.0 NHP long probe (384 recording channels) mounted on a motorized micromanipulator (Narishige) and inserted through a biocompatible cranial implant. We chose recording sites from a square grid of predefined locations spaced 1 mm apart. In each recording session, a single grid location was used, and penetration sites were chosen across sessions to maximize spatial coverage of the DMFC. The recording sites for both subjects are reported in ^56^.

### Spike sorting

Extracellular signals were collected using the spikeGLX software (https://billkarsh.github.io/SpikeGLX), bandpass filtered (300 Hz to 6 kHz) and digitized (sampling rate: 30 kHz). Drift correction of local field potentials (LFP) was applied to the raw signals before spike sorting ^57^. Spike sorting and curation were carried out using Kilosort 2 (https://github.com/MouseLand/Kilosort), followed by manual curation in phy (https://github.com/cortex-lab/phy). Spike trains were binned at 33 ms (30 bins/s), and firing rates were smoothed by convolving spike counts with a Gaussian kernel (σ = 50 ms). For all state-space analyses, each unit’s firing rate was z-scored before principal component analysis (PCA), using the mean and standard deviation of its condition-averaged firing rate.

### Single unit selectivity

We classified single units as start-selective, flow-selective, both, or non-selective using one-way and two-way ANOVAs on single-trial firing rates during three task epochs. Only invisible trials were used. For each unit, firing rates were computed in three windows: an inter-trial interval (ITI) baseline (2.0 to 0.5 s before start-cue onset), a start-position epoch (0.1 s after start-cue onset to flow onset), and a flow epoch (flow onset to 333 ms after flow onset), where flow onset was defined as time zero and start-cue onset occurred 833 ms earlier.

Three analyses-of-variance (ANOVAs) were performed per unit:

1. ITI noise screen: one-way ANOVA of ITI firing rate on start position. Units with p < 0.05 were excluded from selectivity classification to remove units with unstable baselines.
2. Start selectivity: one-way ANOVA of start-epoch firing rate on start position (p < 0.05).
3. Flow selectivity: two-way ANOVA of flow-epoch firing rate on start position and flow direction, with the p-value taken from the direction main effect (p < 0.05).

A unit was classified as start-selective if it passed the ITI screen (p_ITI > 0.05) and had significant start-position selectivity (p_start < 0.05), and as flow-selective if it passed the ITI screen and had significant flow-direction selectivity (p_flow < 0.05). Jointly selective units were both start- and flow-selective; non-selective units were neither start-nor flow-selective.

### Decoding axes for undershoot/overshoot trials

We performed the following analyses on simultaneously recorded populations for each session in Set 2b and Set 3 (animals produced systematic undershoot/overshoot behaviors), and reported results across sessions. Sessions were included if elapsed time during the first 1.0 s of Flow could be decoded from single-trial population activity (cross-validated ridge regression, R² > 0.2; Subj. F, 7 of 12 sessions for Set 2b, 10 of 10 sessions for Set 3; Subj. N, 11 of 13 sessions for Set 2b, 14 of 14 sessions for Set 3).

For each recording session, single-trial neural population trajectories were projected into a PCA space retaining 80% of the total variance. Analysis was restricted to conditions with magnitude-3. Each trial was labelled as "undershoot" or "overshoot" based on its saccade length projection onto the Flow direction: < 3 as undershoot, and ≥ 3 as overshoot. Conditions with fewer than 2 trials in either class were excluded. For each trial, we averaged the neural states of the last five time bins in the Flow period as the neural state for the following decoding analyses.

To test whether the neural encoding of saccade length shared a common axis across Flow directions, we performed three complementary decoding analyses: condition-pooled ("shared") decoder, a condition-specific ("independent") decoder, and a shared-axis-removed (“residual”) decoder.

Shared decoder. A single linear discriminant analysis (LDA) classifier (shrinkage-regularized, solver: least-squares) was trained on all trials pooled across Flow directions, with undershoot/overshoot as the binary label. Classification accuracy was evaluated using leave-one-pair-out cross-validation: on each fold, one (undershoot, overshoot) pair from the same condition was held out, the LDA was trained on all remaining trials. Accuracy was the fraction of correctly cross-validated pairs.

Independent decoder. For each condition separately, LDA was trained on only that condition’s trials using the same leave-one-pair-out procedure. Accuracy was pooled across all conditions.

Residual decoder. (1) a common LDA was fit on all pooled trials (as Shared decoder) to obtain the common discriminant axis; (2) all data were deflated by projecting out this axis; (3) within each condition, an independent LDA (as Independent decoder) was fit on the deflated data using leave-one-pair-out cross-validation. If undershoot/overshoot information is fully captured by the common axis, accuracy after removal should fall to chance. We performed the analyses above for each recording session and reported the results across sessions.

### KiNeT analysis

We adopted the kinematics analysis of neural trajectories (KiNeT) originally introduced in ^18^.

Mean trajectory. For each recording session, we computed a mean trajectory as the trial-averaged neural population trajectory (in PCA space retaining 80% of variance) of the four magnitude-6 conditions. This mean trajectory spans the full displacement duration (2 s for magnitude-6) and captures the population dynamics during the flow epoch covering all shorter-duration conditions.

Undershoot/overshoot trajectories. Individual trial trajectories were extracted from magnitude-3 conditions (Flow duration 1s). Trials were pooled across all magnitude-3 conditions within a session and split into undershoot and overshoot (same criteria as in **Decoding axes for undershoot/overshoot trials**).

Nearest-neighbor time matching. Let *r*(*t*) denote the mean trajectory in PCA space (indexed by time *t*∈[0, *T*]) and *r*’(*t*) denote the group-mean trial trajectory (undershoot or overshoot, indexed by trial time *t*∈[0, *T*]). For each time point *t* on the mean trajectory (T_mean_ in Figs. 2g,i and 3f), we found the time point on the group-mean trajectory whose neural state was nearest in Euclidean distance:

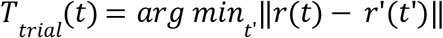

This produces a curve *T_trial_*(*t*) for each group. If a group’s trajectory tracks the mean trajectory at the same pace, *T_trial_*(*t*) = *t* (the identity line). If a group’s trajectory progresses more slowly through the same sequence of neural states, *T_trial_*(*t*) > *t*; *T_trial_*(*t*) < *t* if the group’s trajectory progresses more quickly.

We performed the above analyses for each session with simultaneously recorded neurons, and plotted the s.e.m. across sessions for the undershoot/overshoot groups. At each time bin, the over- and undershoot curves were compared with a one-sided session-stratified permutation test (1,000 permutations), FDR-corrected across bins. The same sessions as for the decoding analysis were used (elapsed-time decoding R² > 0.2).

### Factorization score (ΔR²)

To quantify whether neural population representations factorize task variables (Start and Flow) into additive components, we computed a factorization score based on multivariate linear regression in PCA space.

At each time bin, condition-averaged PC coordinates served as the embedding matrix *E*∈*R^C^*^×d^, where *C* is the number of conditions and *d* the number of retained PCs. The predictor matrix 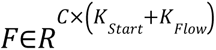 was constructed by concatenating one-hot encodings of Start position (*K_Start_* unique positions) and Flow direction (*K_Flow_* unique vectors). A linear regression without intercept was fit:

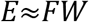

and the coefficient of determination *R*^2^ was computed as the multivariate *R*^2^ across all PCs jointly (scikit-learn r2_score).

To establish a baseline *R*^2^ without additive structure, we permuted the Flow labels within each Start position: the Flow one-hot vectors assigned to conditions sharing that Start were randomly shuffled, breaking any additive direction contribution while preserving marginal start-position structure. The resulting shuffle *R*^2^ reflects variance explained by Start position alone plus any spurious direction fit from the permuted labels.

The factorization score was defined as 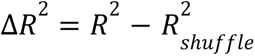. A positive Δ*R*^2^ indicates that the population geometry contains additive structure more than the spurious direction fit to start alone. We ran the analyses for each recording session (using only Start locations paired with more than one Flow direction) and presented the results across sessions.

### Correlating neural deviation with behavioral error

To test whether the departure of factorized Flow trajectories from a well-learned template predicts behavioral error, we performed the following analysis.

For each animal, condition-averaged firing rates from the last five Set 2a sessions were pooled across sessions into a pseudo-population, forming a firing-rate tensor *R*∈*R^N^*^×*C*×*T*^, where *N* is the total number of pseudo-units, *C* the number of conditions, and *T* the number of time bins. Only invisible trials were included. The tensor was projected into a PCA space retaining 80% of the total variance.

We identified the eight conditions in Set 2a that formed clockwise and counter-clockwise circles around the fixation point (magnitude = 2; two conditions per Flow direction), which exhibited the lowest behavioral error across both subjects. We designated these as template conditions and hypothesized that their Flow-epoch population trajectories (template trajectories) are essential for successful behavior in other conditions.

For each non-template condition, we computed the Euclidean distance to each same-direction template trajectory in PCA space at each time bin (two templates per Flow direction). All trajectories were first baseline-subtracted by their value at Flow onset (*t* = 0), so the distance reflects divergence in the flow-evoked response:

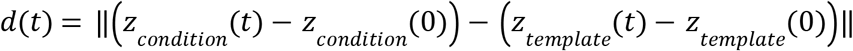

where *z*(*t*) is the PCA-space trajectory at time *t*.

The analysis was evaluated at the last time bin of the Flow epoch. Each condition-to-template pair yielded one neural distance value and one associated behavioral error (the normalized saccade error of the non-template condition). Both quantities were *z*-scored across condition pairs, and their relationship was assessed using Pearson correlation.

### Kernel-based subspace alignment

To compare neural population trajectories across separately recorded pseudo-populations (Set 2a and Set 3 subsets), we developed a kernel-based alignment procedure that projects each pseudo-population into a shared low-dimensional subspace *K*, guided by a similarity metric from the **Gaussian Process Model of behavior**.

For each animal and condition set (Set 2a or Set 3 subsets), condition-averaged firing rates from all sessions were pooled into a pseudo-population tensor (*N* units × *C* conditions × *T* time bins; only invisible trials). Each pseudo-population was independently PCA-reduced, retaining components explaining ≥ 80% of variance. All time bins were then projected through the retained basis, yielding per-set PCA trajectories of dimension *k_set_* and *k_gen_* (which may differ).

A target pairwise distance matrix *d_kernel_*(*i*, *j*) was computed over all conditions from both sets using the Matérn-5/2 kernel from the behavioral GP fit (see **Gaussian Process Model of behavior**). The kernel had unit amplitude and used the GP lengthscales (geometric mean of the *x* and *y* fits), so that 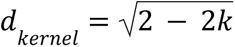.

For each set, a linear projection matrix mapped the PCA trajectories into a shared *K* dimensional subspace. We used *K* = 3 for Fig. 3j (Fig. S3.2 shows *K* = 3 − 10), and neural deviation for Fig. 3j was measured 0.33 s after Flow onset. The projected trajectory of condition *c* at time *t* is:

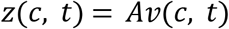

where *v* is the PCA-space trajectory and *A* is the set-specific projection matrix (*A_set_* or *A_gen_*). The neural distance between conditions *i* and *j* was the Euclidean distance in the shared subspace, averaged over a time window restricted to the flow epoch common to both conditions:

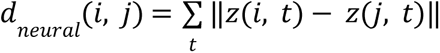, where the average is taken over *t*∈[0, *min*(*dur_i_*, *dur_j_*)], the flow epoch shared by both conditions.

The projection matrices were optimized jointly to minimize the mean squared error between neural and kernel distances:

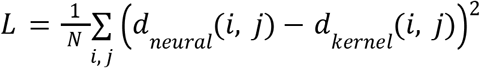

*A_set_* and *A_gen_* were optimized jointly using torch.optim.Adam (PyTorch).

### Recurrent neural networks

#### Input format

Each trial was represented as a time series of 81 bins (bin width 33.3 ms, total duration 2.7 s), structured as four epochs: fixation (0–0.3 s, 9 bins), flow (0.3 s onward, duration determined by displacement magnitude at Flow speed 3 grid units/s, ranging from 0.33 s for magnitude 1 to 2.0 s for magnitude 6), delay (0.2 s, 6 bins), and report (0.2 s, 6 bins). The model received two input streams at each time step. The Flow input was a 4-dimensional one-hot encoding of direction, active only during the flow epoch and zero otherwise. The Start input was a 49-dimensional retinotopic population code in which each channel corresponded to one node of the 7×7 grid (centers uniformly spaced from −3 to 3 in both *x* and *y*) and was computed as a Gaussian bump, 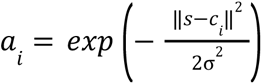, where *s* is the start position, *c*_i_ the center of the *i*-th node, and σ = 1 grid unit. The Start code was tonic (constant across all time steps within a trial). The output target was the 2-dimensional endpoint (x, y), active only during the report epoch; all other time bins were masked (NaN) and excluded from the loss.

#### Recurrent dynamics

All recurrent units followed a continuous-time formulation discretized with step size Δ*t* = 1/30 s and time constant τ = 0. 1 s, giving a leak rate 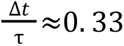:

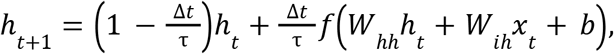

where *f* was the *tan*ℎ nonlinearity unless otherwise specified. All recurrent networks had hidden size 32.

#### Training protocol

Each model was trained on the same conditions used to train the animals (Sets 1 and 2, 108 conditions) and evaluated on all remaining valid conditions. On each trial, the RNN processed the full input time series and its hidden state was read out at the report epoch to produce a single 2-dimensional endpoint prediction. The primary loss was the mean squared error (MSE) between the predicted and true endpoint. Auxiliary losses were imposed for different model variations. Training used the AdamW optimizer (initial learning rate 10 ⁻ ³, no weight decay, gradient clipping at 1.0). Each architecture was trained 10 times with different random initializations; results were pooled across seeds.

#### Timing Module pre-training

The Timing Module is an RNN pre-trained on a pure timing task before being incorporated into the full architecture. In this pre-training stage, the RNN received a 1-dimensional step input of constant amplitude for a variable duration and learned to produce, through a linear readout, a linear ramp from 0 to the flow duration in seconds. The six durations matched the task (magnitude 1–6 at Flow speed 3 grid units/s, yielding 0.33–2.0 s). This stage used AdamW (learning rate 3 × 10 ⁻ ³, gradient clipping at 1.0), and converged when MSE fell below 10 ⁻ ³. After convergence, the recurrent weights (*W_hh_* and bias) and the ramp readout *w_ramp_* were frozen. The input weights *W_i_*_ℎ_ were re-initialized to accept the 4-dimensional Flow signal and remained trainable for subsequent tasks.

#### Model a (Fig. 4a)

A full-rank RNN received the Flow input and was trained end to end without pre-training. At flow offset, the hidden state was concatenated with the Start population code and mapped to the target through a linear layer (no nonlinear readout). The loss was purely behavioral (no auxiliary supervision).

#### Model b (Fig. 4b)

The frozen Timing Module received the Flow input through trainable input weights. At flow offset, the hidden state was concatenated with the Start population code and passed through a linear network to produce the target prediction. We imposed an auxiliary loss *L_ramp_*, defined as the MSE between the frozen ramp readout *w_ramp_* applied to the hidden state and an ideal linear ramp. This loss ensured that the trainable input weights preserved the temporal integration structure established during pre-training. The total loss was *L* = *L_be_*_ℎ*av*_ + 30*L_ramp_*.

#### Model c (Fig. 4c)

Same as **Model b**, but hidden state was concatenated with the Start population code and passed through a feedforward network (six hidden layers of 128 units each with tanh activation) to produce the target prediction.

#### Model d (Fig. 4d)

Same as **Model c**, except that a nonlinear embedding jointly transformed the two input streams before they entered the architecture. At each time step, the Flow (4D) and Start (49D) were concatenated into a 53-dimensional vector and passed through a feedforward network with two hidden layers of 32 units ([53, 32, 32, 53], tanh). The output was split back into a 4-dimensional flow signal (routed to the frozen Timing Module) and a 49-dimensional start signal (concatenated with the hidden state at the readout).

### Evaluation of the three constraints

(1) Shared-axis constraint. We performed two analyses. First, a PCA visualization on the raw trajectories and a residual PCA (insets) after subtracting the per-time-step mean across directions (Start state removed) to highlight direction-specific divergence. The same analysis was applied separately to four magnitude-6 conditions (one per direction) from the training set and four from the holdout set (Fig. 4). Second, a perturbation experiment to evaluate the causal relationship between integration axes and produced displacement length. The common axis is the direction of the mean trajectory velocity averaged across all four cardinal flow directions. The residual axis is the deviation of a single direction’s endpoint from the mean endpoint; it captures direction-specific information caused by the recurrent nonlinearity. For each holdout condition, the hidden state ℎ at flow offset is perturbed by ±ε (ε = 0. 2) along each axis, the output displacement length is recomputed through the readout, and the signed slope 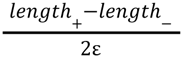 quantifies how sensitively the readout maps movement along that axis onto predicted magnitude (Fig. S4.2).
(2) Error-coupling constraint. For each holdout condition, we identified the same-direction, same-magnitude training condition with the lowest normalized error as the template. We removed the Start state of each trajectory and computed the neural deviation as the Euclidean distance between the holdout and template trajectories at the last time bin of the Flow epoch, as in the monkey analysis. The neural deviation was then correlated with the normalized behavioral error of the holdout condition. Neural deviation was z-scored within magnitude; Pearson correlation with behavioral error was computed across conditions pooled over seeds.
(3) Proximity constraint. For each holdout condition, we computed the minimum Euclidean distance in 4-d condition space (*x*_0_, *y*_0_, Δ*x*, Δ*y)* to training conditions sharing the same direction. Both distance and behavioral error were z-scored within magnitude, and their Pearson correlation was computed across conditions, with error averaged across seeds.

#### Full generalization model (Fig. 5f)

Identical to Model c, except that the readout was restructured to separate the computation of displacement from its combination with Start. The hidden state at flow offset was passed through an MLP (32 → 32 → 32 → 2, tanh) supervised to output the true displacement vector in 2D Cartesian space. The displacement loss (*L_disp_*) was the MSE between the displacement MLP output and the true displacement vector *d*×*t*, computed at flow offset. The Start population code (49D) bypassed the MLP and was concatenated with the predicted displacement, and a single linear layer (51 → 2) produced the target. The total loss was *L* = *L_be_*_ℎ*av*_ + *L_disp_* + 30*L_ramp_*.

#### Two-dimensional parametric sweep (Fig. 5g)

To characterize the contributions of factorized input and task-aligned output to generalization, we varied the **full generalization model** along two axes toward an entangled, general-readout variant. This variant resembles Model d (Fig. 4d) but keeps the displacement MLP and uses a smaller readout network (see below).

The first parameter α controls the degree of input factorization. At α = 1, only Flow entered the embedding and Start reached the readout directly; at α = 0, Start and Flow were jointly transformed by the embedding. Flow passed through the embedding at all values of α. To interpolate between these extremes, the embedding received [*Flow*(4*D*); (1 − α)·*Start*(49*D*)], and the Start representation at the readout was blended as *start_readout_* = (1 − α)*start_mix_* + α*start_direct_*, where *start_mix_* was a linear readout (32 → 2) of Start from the Timing Module hidden state and *start_direct_* was the linear projection (49 → 2) of the Start population code. We swept α from 0 (fully entangled) to 1 (fully factorized).

The second parameter β controls the degree of task alignment in the output. In the full generalization model, the predicted displacement and *start_readout_* were combined through a single linear layer ([2D; 49D] → 2D), which matches the additive structure of the vector composition task. In Model d, the combination was a six-hidden-layer FFN with scaled tanh activation, which can learn arbitrary input–output mappings without built-in task structure. To interpolate between these extremes, we replaced the linear readout of the full generalization model with a two-hidden-layer FFN (51 → 32 → 32 → 2) using the scaled tanh activation 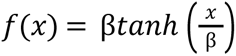. When β is small, the activation saturates sharply and the network can fit arbitrary nonlinear mappings; as β→∞, the activation approaches the identity and the FFN reduces to a linear transformation.

The sweep covered 25 conditions:

α∈{0, 0. 25, 0. 5, 0. 75, 1. 0}×β∈{0. 125, 0. 5, 1, 4, ∞(*linear*)}

### Human behavioral experiments

#### Participants

49 participants were recruited through the Prolific platform (https://www.prolific.com). The protocol was approved by the Committee on the Use of Humans as Experimental Subjects at the Massachusetts Institute of Technology. Participants gave informed consent before the experiment. Sessions lasted a median of 27 min and participants were compensated $12 (approximately $27/hour at the median session length). Participants could withdraw at any time. The participants (28 male, 21 female; self-reported sex from Prolific) were 19–67 years old (mean 32.2 ± 10.6 s.d.).

#### Task structure

Participants performed a target-localization task on a black 800 × 800 pixel canvas displayed in a web browser. A white fixation cross was presented at the canvas center (400, 400). Each trial began with a 1,500 ms inter-trial interval, after which the Start location appeared as a yellow circle (radius 20 px) at a designated position on the canvas. After 1000 ms, a Flow stimulus was presented around the center of the screen. The Flow consisted of coherent moving dots moving at a speed of 100 px/s. In Experiments 1 and 2, the dots translated in one of four cardinal directions (Cartesian flow). In Experiment 3, the dots rotated or expanded/contracted within the aperture (polar flow), matching the direction and speed of the polar target motion. The duration of the Flow determined the displacement magnitude, similar to the monkey experiment. Following Flow offset, participants clicked on the canvas to indicate the inferred target location. Only the first click was registered. The correct target was defined by the vector composition of the Start location and the Flow-specified displacement, computed in Cartesian coordinates for Experiment 1 and in polar coordinates relative to the screen center for Experiments 2 and 3. A success was registered when the Euclidean distance between the click and the target fell within a tolerance window that scaled with displacement magnitude. After a click, a coin animation appeared on the target. A success sound indicated a correct response, and a buzzer indicated an error. For each experiment, subjects received visible training trials at a small number of start positions and were then tested on holdout positions without visual guidance. In visible trials, a gold coin traced the displacement from the Start toward the Target during the Flow period, allowing participants to solve the task by tracking the moving cue. In invisible trials, no visual guidance was provided; participants relied entirely on the learned mapping to infer the target location.

#### Experiment 1 (Cartesian Flow → Cartesian motion)

At the beginning of the experiment, subjects were told to infer the underlying rule of the game without explicit instruction. Subjects were first presented with 12 visible conditions as training. The conditions spanned 2 start positions ([−100, −100] and [100, 100] px relative to center), each with 2 of the 4 cardinal directions (together covering all four), and 3 durations (1, 2, 3 s). The integrated displacement of the flow corresponds to the displacement of the target from the start, creating a congruent mapping. Subjects were then tested on 36 holdout conditions, where only invisible trials were presented. Those holdout conditions contained start locations not included in the training set, evenly distributed across the canvas.

#### Experiment 2 (Cartesian Flow → Polar motion)

After Experiment 1, subjects were informed that the underlying rule would change in the next stage. In Experiment 2, the same Cartesian flow stimulus was mapped to polar displacements relative to the screen center: upward flow → radial expansion, downward flow → radial contraction, leftward flow → counter-clockwise rotation, rightward flow → clockwise rotation.

Subjects were trained at 2 start positions near the bottom of the screen ([−50, −150] and [50, −200] px relative to center), where the polar motion was incompatible with the intuitive understanding of Cartesian flow. The training set comprised 78 trials (40 visible, 38 invisible) jointly covering all 4 directions and matched durations (rotation: 1, 2, 3 s; radial: 0.75, 1.25 s).

Subjects were then tested on holdout conditions spanning 14 start positions on two concentric rings (radii 150 and 250 px from center) at 7 equally spaced angles covering ±120° from the bottom of the screen. The top 120° was excluded because the polar and Cartesian mappings are approximately congruent there. Each start position was tested with a balanced set of directions and durations (96 probe trials). Holdout trials were presented in an interleaved order that decorrelated angular distance from trial index, controlling for potential confounds of degraded performance across time.

#### Experiment 3 (Polar → Polar)

After Experiment 2, subjects were informed once more about the change of task rules. Subjects were then presented with polar Flow that indicated polar motion of the target. For radial Flows (expand, contract), the dot velocity tracked the target’s velocity along the radius; for rotational Flows (CW, CCW), the dots’ angular velocity tracked the angular velocity of the target relative to screen center, creating a congruent and intuitive mapping between stimulus and displacement.

Subjects were trained on 8 visible conditions at 4 start positions at angles −120°, −40°, 40°, 120° from the bottom of the screen, balanced across Flow type and durations exposed in Experiment 2. Subjects were then tested on 28 holdout conditions without visible trials (14 Starts covering ±120° × 2 conditions each, 1 rotational + 1 radial, balanced across types and durations).

## ACKNOWLEDGEMENTS

We thank Mahdi Ramadan and Jack Gabel for their guidance in setting up the experimental rigs and supporting our neurophysiology experiments. We thank the CatalystNeuro team for their help open-sourcing our data. We thank Srdjan Ostojic, Arianna Bernardo, and members of the Jazayeri lab for their feedback on the findings.

## AUTHOR CONTRIBUTIONS

C.T. and M.J. designed the task. C.T. collected the data. C.T. and M.J. developed the analyses and models. C.T. did the data processing and model implementation. C.T. and M.J. wrote the manuscript.

## FUNDING

M.J. is supported by SCGB, HHMI, and the McGovern Institute.

## COMPETING INTERESTS

The authors declare no competing interests.

## DATA AND CODE AVAILABILITY

The data used to generate all results will be made available on a public repository after peer-review publication.

The code used to generate all results will be made available on a public repository after peer-review publication. The code used for all the models is available on a public repository: https://github.com/jazlab/CT_MJ_2026

## Supplementary Figures

**Supplementary Figure S1.1.**
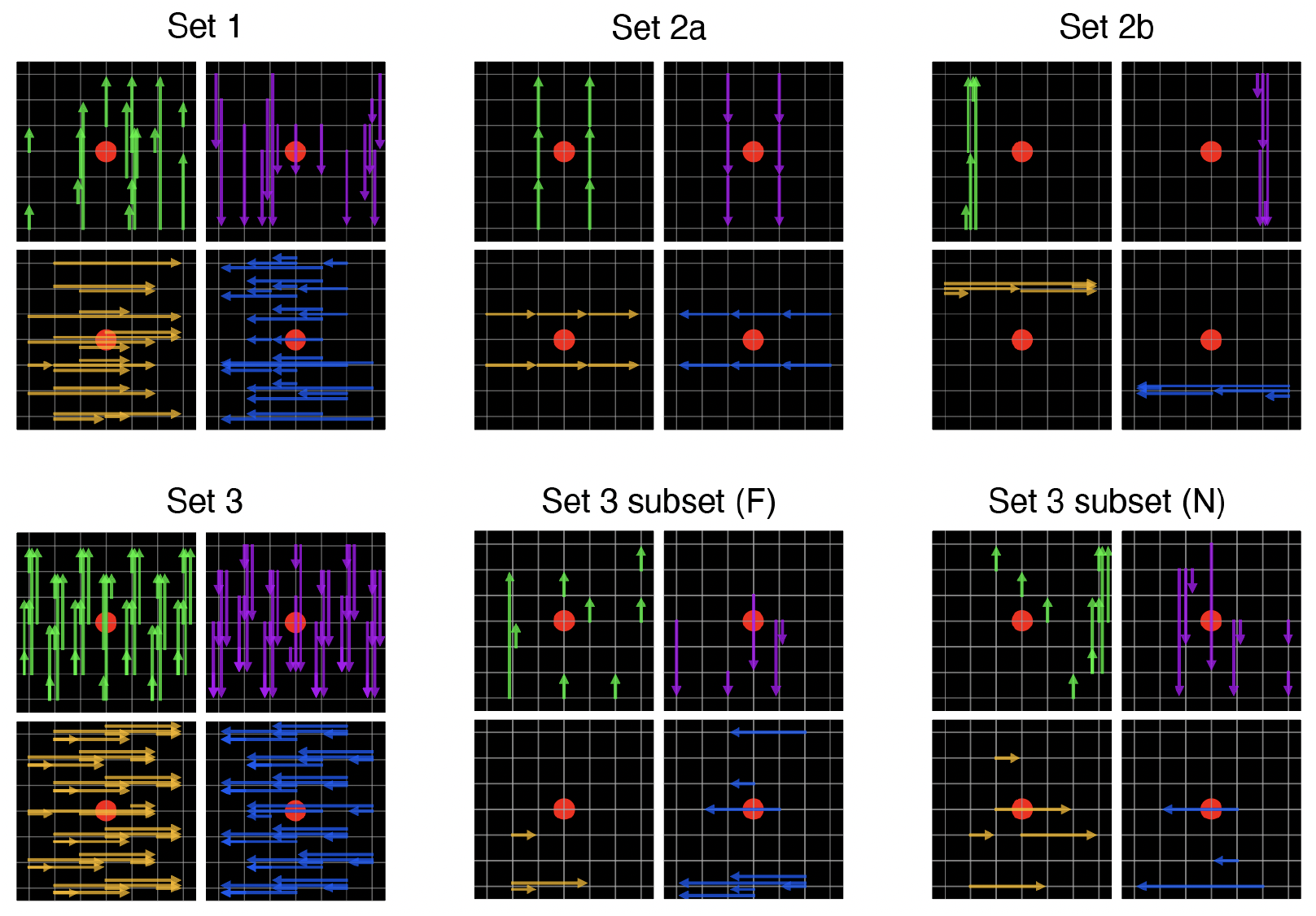
Complete experimental conditions across all sets. Set 1: 74 conditions. Set 2a: 24 conditions. Set 2b: 20 conditions. Set 3: 129 holdout conditions. Set 3 subsets: 24 conditions for Subj. F, 26 conditions for Subj. N. Arrows point from Start to Target.

**Supplementary Figure S1.2.**
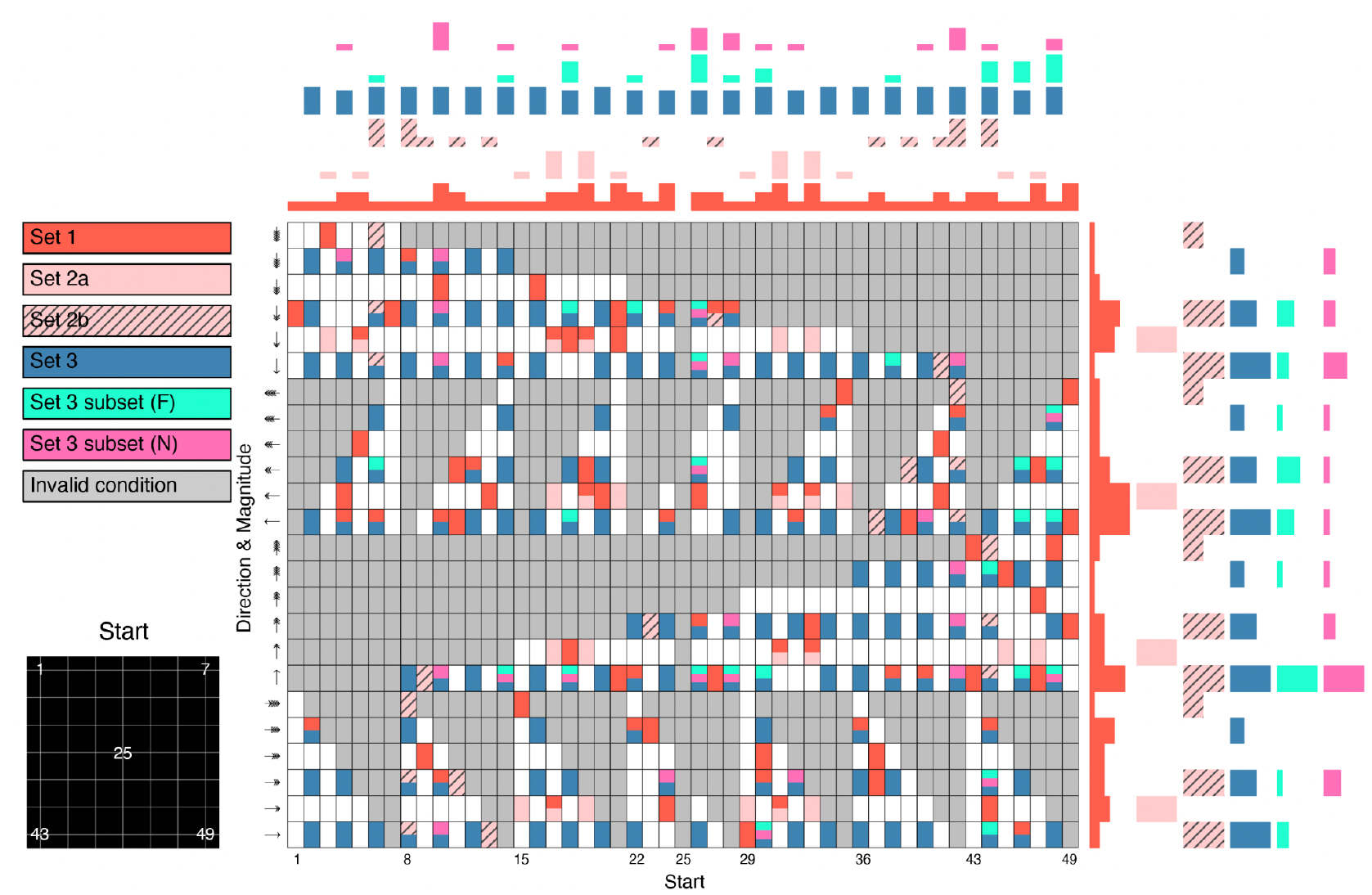
Start–Flow matrix across all experimental sets. Each cell in the matrix represents a unique Start–Flow pairing. The x axis indexes the 49 Start locations on the 7×7 grid; the mapping from index to grid position is shown in the inset at bottom left (index 1 = top-left corner, index 25 = center, index 49 = bottom-right corner). The y axis indexes the 24 displacement vectors, grouped by direction (arrows) and ordered by increasing magnitude within each direction. Cell colors indicate set membership. Gray cells denote invalid conditions. Marginal histograms along the top and right edges show the number of conditions from each set. Conditions in Set 1 were designed to span all Start and Flow options independently (marginal coverage along both axes).

**Supplementary Figure S1.3.**
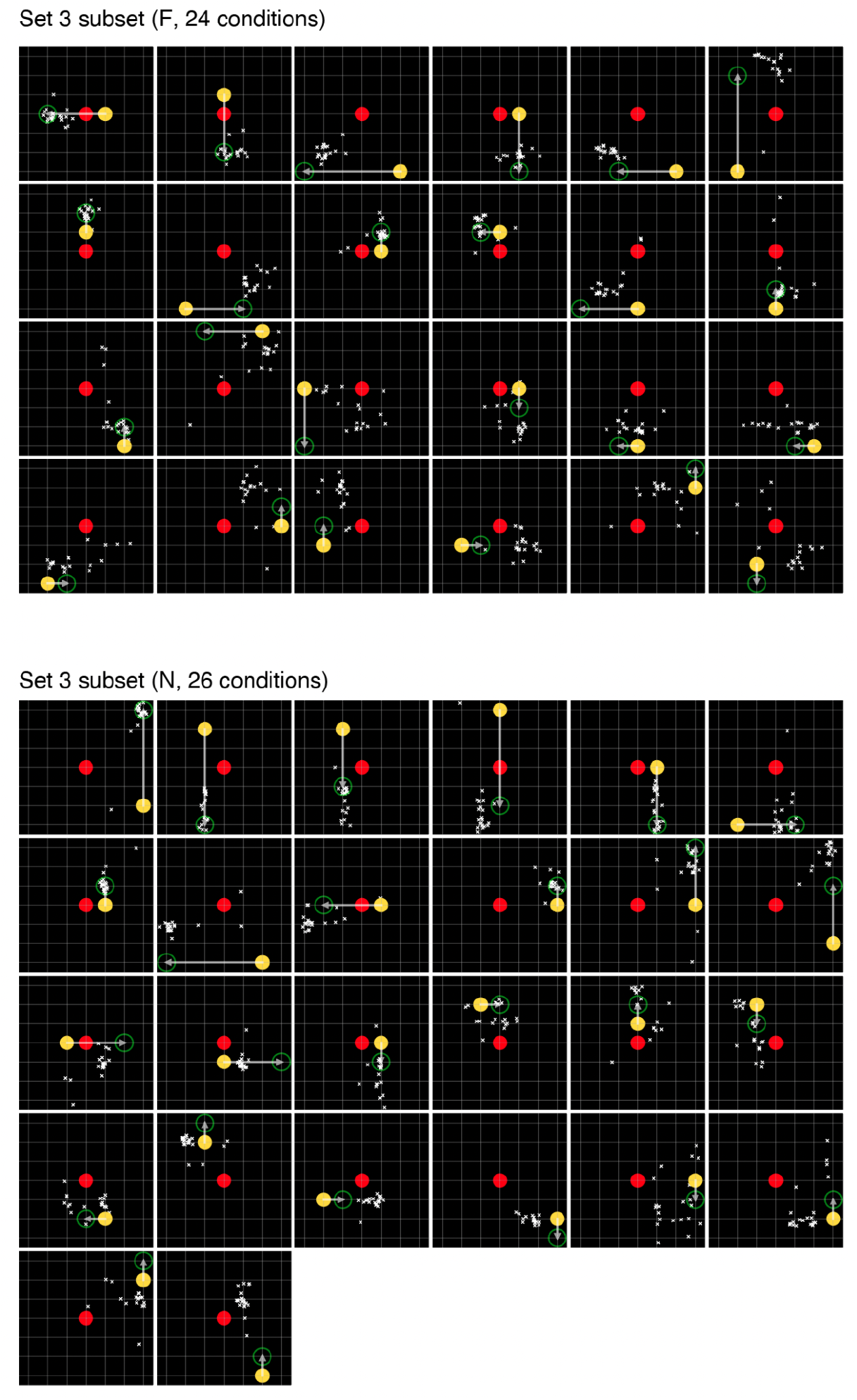
Single-trial behavior for Set 3 conditions. Each panel shows one held-out condition. Subj. F, 24 conditions; Subj. N, 26 conditions. Yellow, Start; red, fixation; green, Target; arrow, Flow vector; white crosses, individual saccade endpoints on invisible trials from the first five Set 3 sessions (*n* ≈ 19–21 trials per condition). We used these behaviors to select a subset of conditions for neural recordings, covering conditions that were either consistently correct or systematically biased in each animal. Panels are ordered by normalized error, from lowest (top left) to highest (bottom right).

**Supplementary Figure S1.4.**
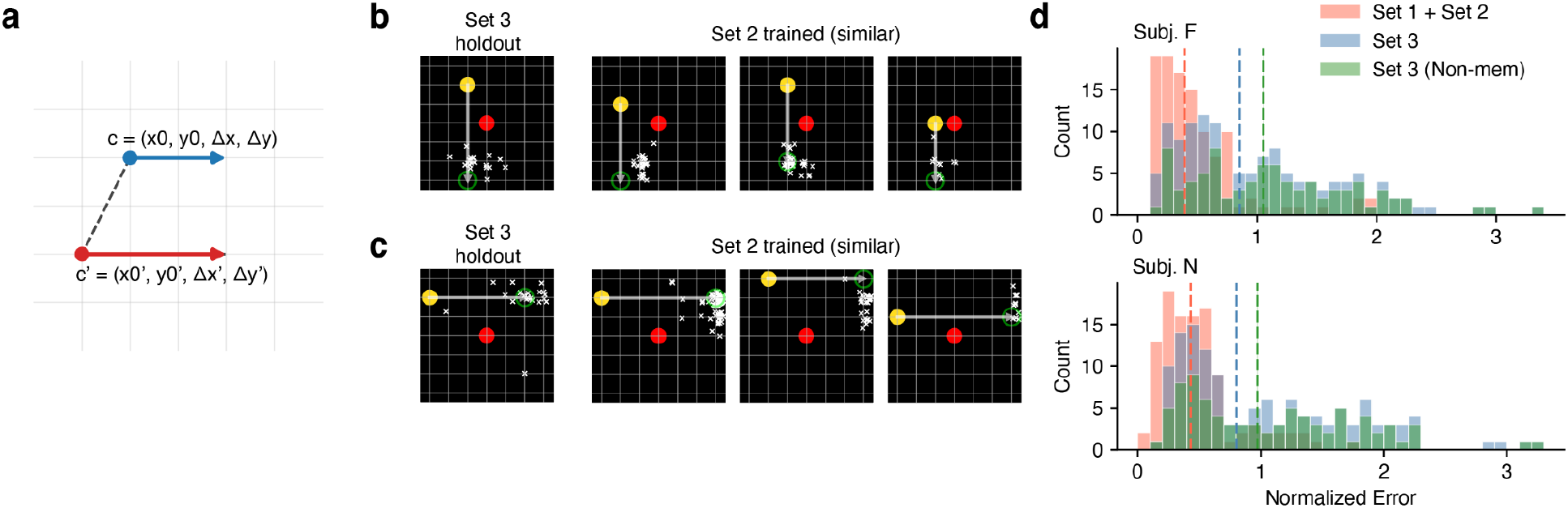
Similarity-based memorization model. **a**, Similarity metric. For each Set 3 condition, we computed the Euclidean distance to every trained (Set 1 + Set 2) condition in the four-dimensional space of **c** = (*x*_0_, *y*_0_, Δ*x*, Δ*y*) – the Start position (*x*_0_, *y*_0_) and the Flow displacement (Δ*x*, Δ*y*) – as *D*^2^ = Σ_i_ (*c*_i_ − *c*_i_′)^2^. To evaluate if the response to a holdout condition is consistent with a memorization strategy, we compared the saccade endpoints against those from the five most similar training conditions (smallest *D*), using energy distance analysis. Energy distance between two point clouds *A* and *B* was computed as: *E*(*A*, *B*) = 2⟨*d*_AB_⟩ − ⟨*d*_AA_⟩ − ⟨*d*_BB_⟩, where ⟨*d*_AB_⟩ denotes the mean pairwise Euclidean distance between points in *A* and *B*, and ⟨*d*_AA_⟩ and ⟨*d*_BB_⟩ denote mean within-set distances. Statistical significance was assessed via permutation test (1,000 iterations). On each iteration, saccade endpoints from both conditions were pooled, randomly relabeled into two surrogate sets matching the original sample sizes, and the energy distance was recomputed to construct a null distribution. The *p*-value was the proportion of permutations yielding an energy distance ≥ the observed value. A hold-out condition was classified as "non-memorization" if its saccade distribution differed significantly (p < 0.05) from all five nearest training conditions. **b–c**, Markers same as S1.3. **b**, An example Set 3 condition plausible with memorization. Left: a Set 3 condition, Right: five conditions in Set 1 + Set 2 with smallest *D* (three plotted). Saccade endpoints from Set 3 condition are not significantly different from those on the right (energy distance permutation, p>0.05). **c**, Same as b, but inconsistent with memorization. The saccade distribution for the Set 3 condition is statistically distinct from all similar trained conditions (energy distance permutation, p < 0.05). **d**, Normalized errors for trained conditions (Set 1 + Set 2, red), holdout conditions (Set 3, blue), and non-memorization Set 3 conditions (green) for each animal. Dashed lines indicate the median of each distribution. Non-memorization conditions (Subj. F: 103/129, 80%; Subj. N: 98/129, 76%) exhibit higher errors than the full Set 3.

**Supplementary Figure S2.1.**
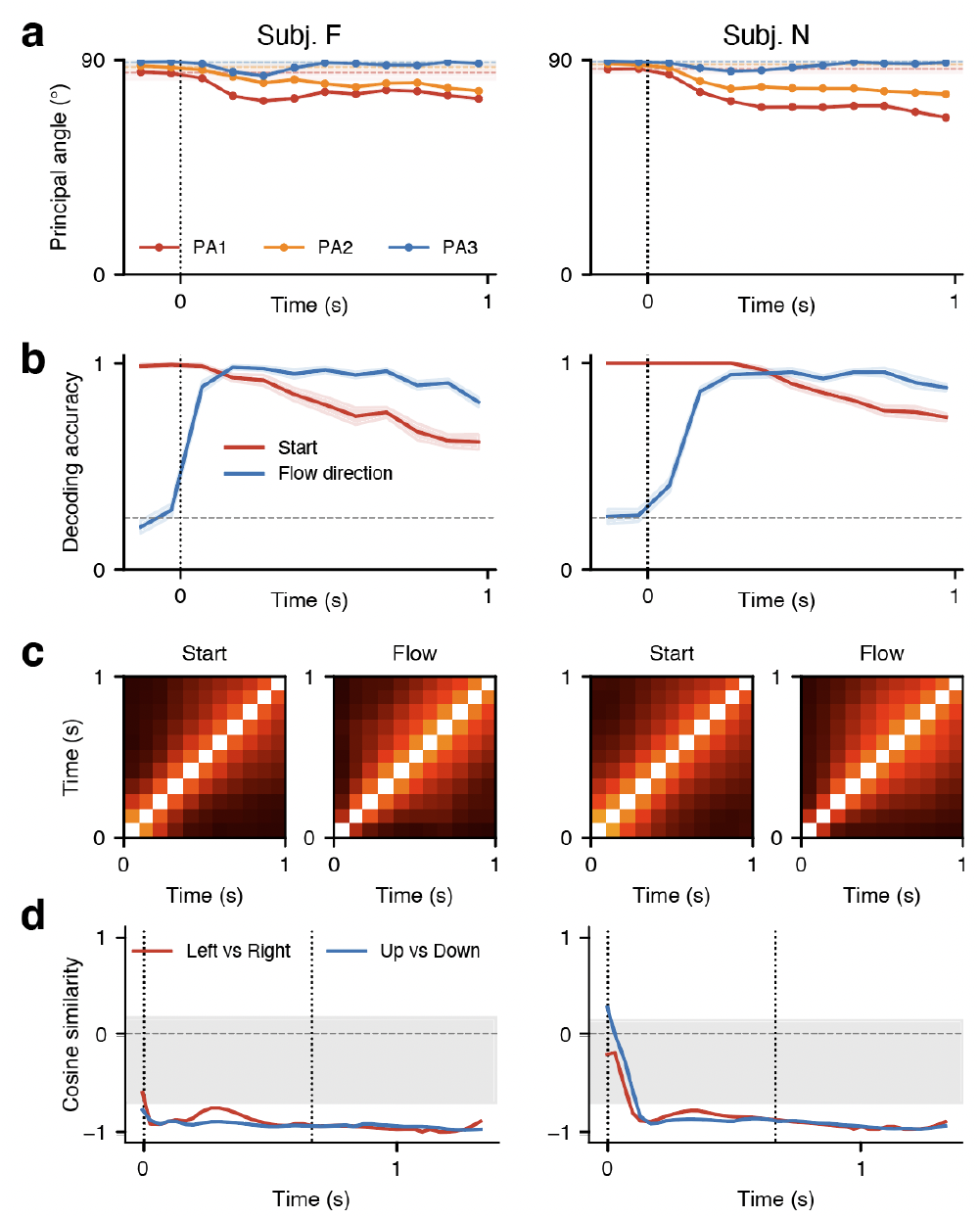
Encoding of Start and Flow information. Left and right columns, Subj. F and N. Analysis uses 16 fully-crossed conditions from Set 2a (4 center Start positions × 4 Flow directions). Start and Flow subspaces are defined by SVD on condition averages grouped by Start or Flow identity. Error bands indicate ± SEM across 10 cross-validation iterations. **a**, Principal angles (PA) between the Start and Flow subspaces over time. PA1, PA2 and PA3 are the three principal angles sorted from most to least aligned. Dashed lines and shaded bands show the null distribution from random subspaces of matching dimensionality. **b**, Cross-validated decoding accuracy over time for start position (red, 4 classes) and flow direction (blue, 4 classes) from the full pseudo-population of Set 2a. Nearest-centroid classifier; dashed gray line indicates chance (1/4 = 0.25). **c**, Cross-temporal subspace alignment for Start (left) and Flow (right) encoding subspaces during the flow period. Each cell shows the mean principal angle between the subspace defined at one time point and one at another. Light cells (low angle) indicate stable subspace orientation; dark cells (high angle) indicate rotation. **d**, Cosine similarity between opposite Flow stimuli. Firing rates (−1.0 to 1.33 s around Flow onset) are projected via global PCA (80% variance; Subj. F: 17D, Subj. N: 18D). We first averaged trajectories per Flow direction. At each time bin, we removed the mean trajectory (centroid) across Flow directions after projecting out the component along the mean trajectory’s tangent. Cosine similarity is then measured between Flow-to-centroid vectors for each opposite pair: left vs right (red) and up vs down (blue). Gray band shows the 95% CI of the cosine for one opposite pair of random direction vectors, centered across the four directions as in the data (Subj. F: 16D, [−0.71, 0.17]; Subj. N: 17D, [−0.70, 0.16]; 10,000 draws; null mean −1/3).

**Supplementary Figure S2.2.**
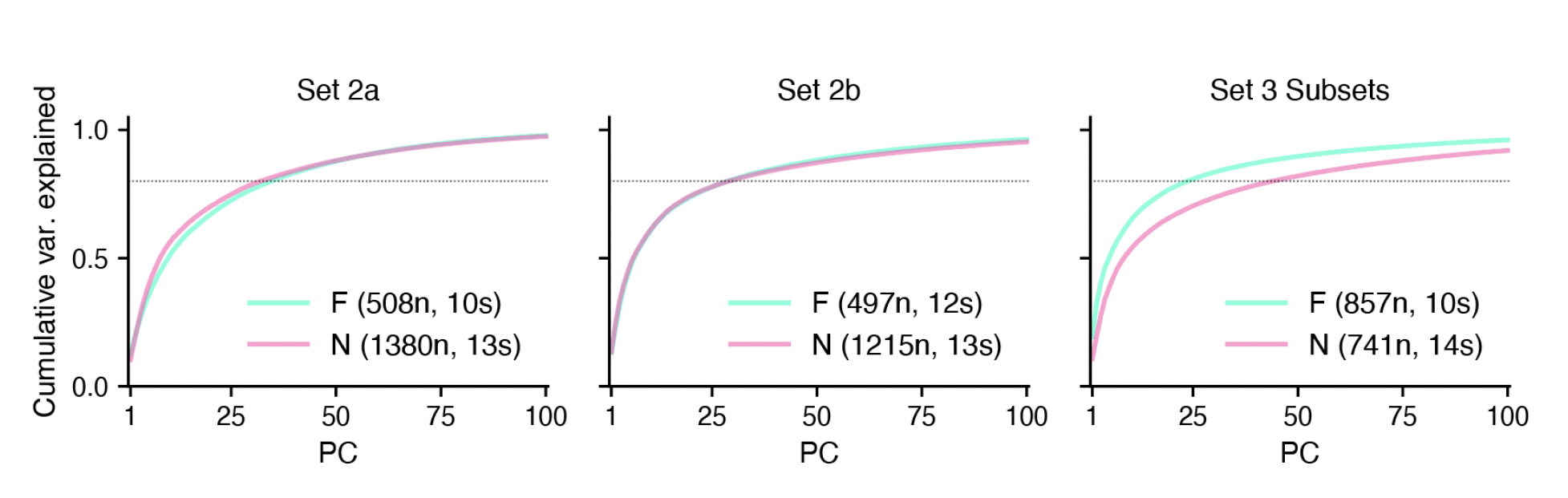
Cumulative variance explained by principal components. Scree plots for the pseudo-populations used in the state-space visualizations (Fig. 2). Each panel shows cumulative variance explained as a function of the number of PCs retained, for Subj. F (cyan) and Subj. N (pink). Pseudo-populations were constructed by concatenating neurons across sessions (*n* = number of neurons, *s* = number of sessions). PCA was fit on condition-averaged, z-scored firing rates during the flow period. The dashed line indicates 80% variance explained. The first 2 PCs capture 18–32% of variance across datasets (Set 2a: 19.1% F, 18.4% N; Set 2b: 23.1% F, 24.8% N; Set 3 Subsets: 32.1% F, 19.4% N); 80% variance is reached by approximately 20–40 PCs depending on the dataset and animal.

**Supplementary Figure S3.1.**
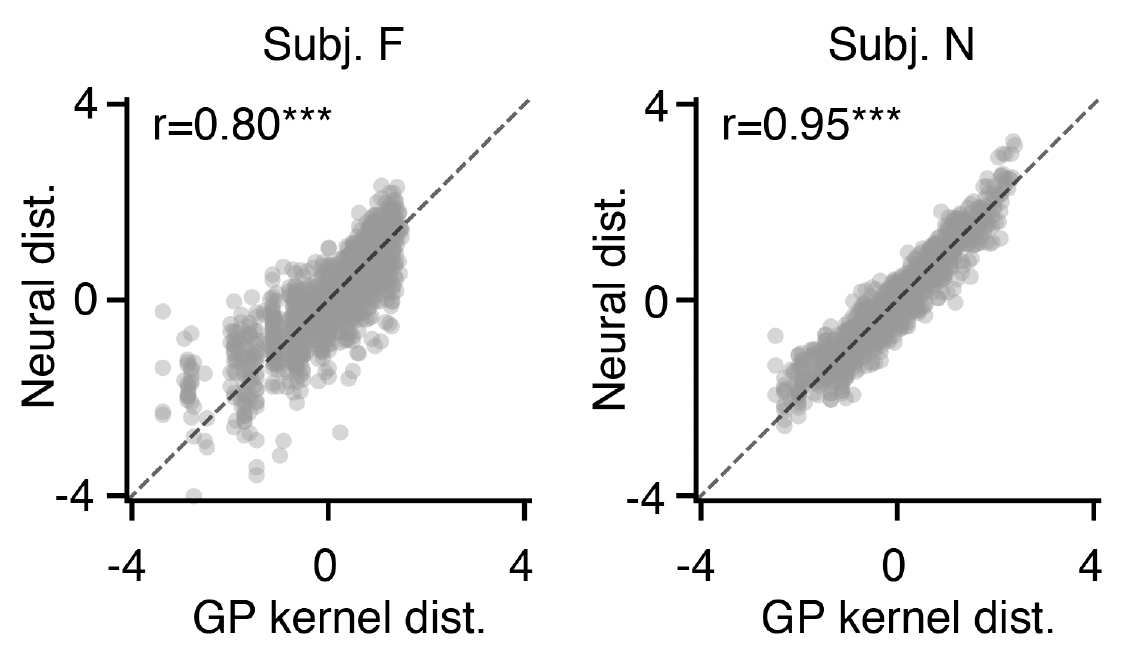
Kernel-based subspace alignment recovers GP kernel distance structure in neural population trajectories. Scatter plots of z-scored kernel distance (x-axis) versus z-scored neural trajectory distance in the aligned subspace (y-axis) for each condition pair. The dashed line indicates identity. The optimization closely recovers the target kernel geometry in both animals (Subj. F: Pearson’s r = 0.80; Subj. N: r = 0.95).

**Supplementary Figure S3.2.**
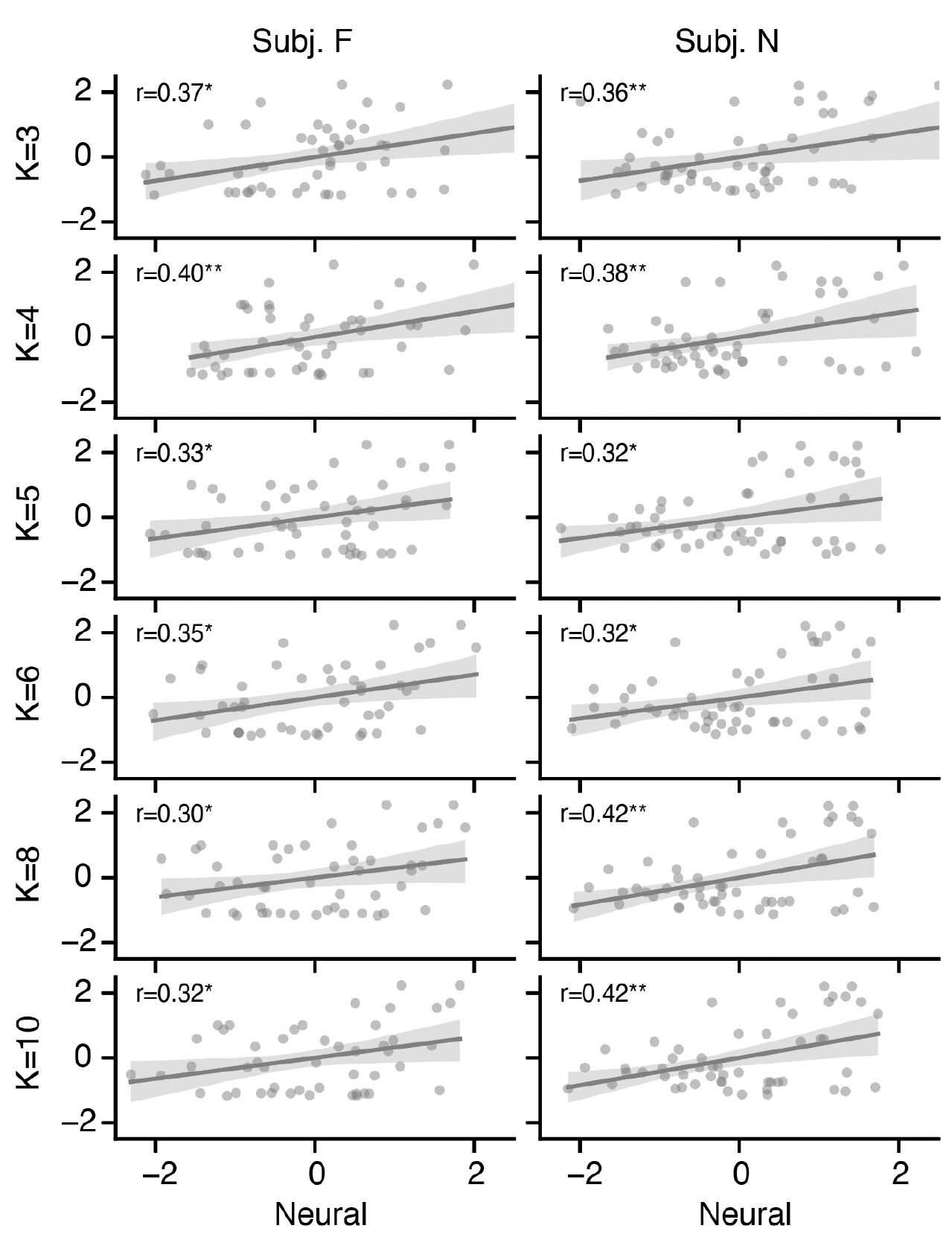
Neural-deviation to behavior-error correlation is robust to shared subspace dimension. **K.** We replicated Fig. 3j for different subspace dimensions K = 3–10. Each column shows one animal (Subj. F, left; Subj. N, right); each row corresponds to a different K. Gray lines show linear regression fits with 95% confidence bands. The correlation is significant (Pearson’s r, * p < 0.05, ** p < 0.01) for all K values in both animals (Subj. F: r = 0.30–0.40; Subj. N: r = 0.32–0.42).

**Supplementary Figure S3.3.**
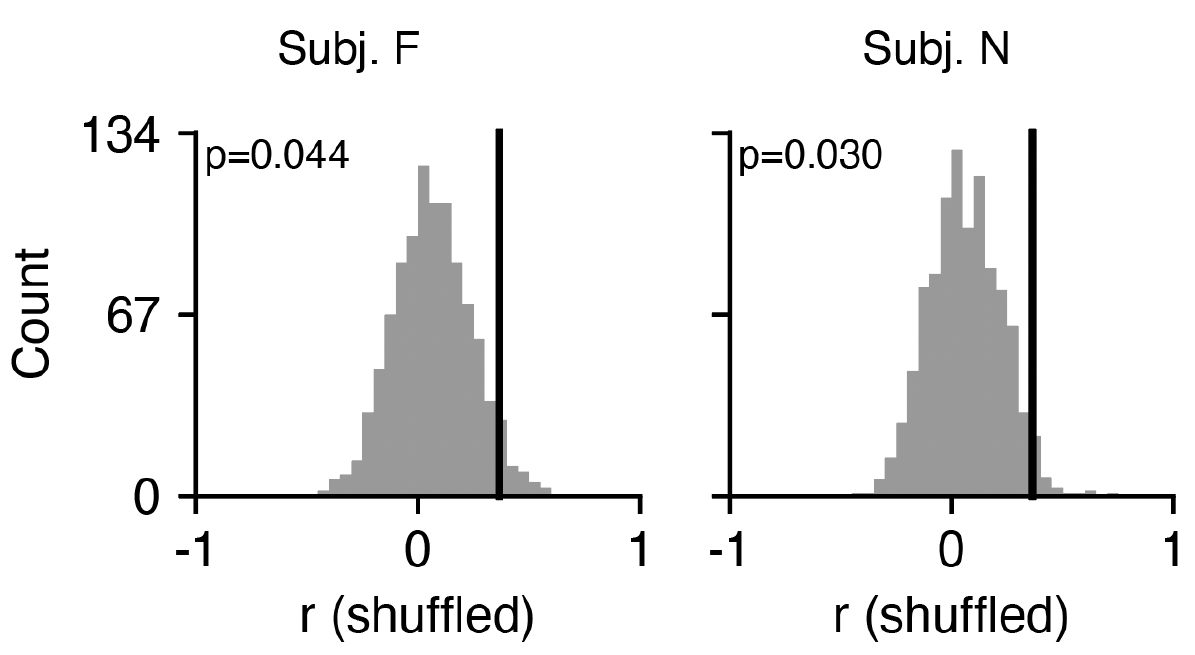
The correlation between neural deviation and behavioral error depends on the geometry of Set 3 trajectories. Distribution of Pearson’s r between neural deviation and behavioral error (computed as in Fig. 3j) after shuffling Set 3 trajectories among conditions with the same Flow magnitude and refitting the kernel-based alignment (1,000 permutations). Each condition’s coordinates, kernel distances, behavioral error and template assignment were kept at their true values; only the neural trajectory was reassigned. The black line marks the observed correlation (Fig. 3j). Permutation p-values are one-sided (Subj. F: p = 0.044; Subj. N: p = 0.030).

**Supplementary Figure S4.1.**
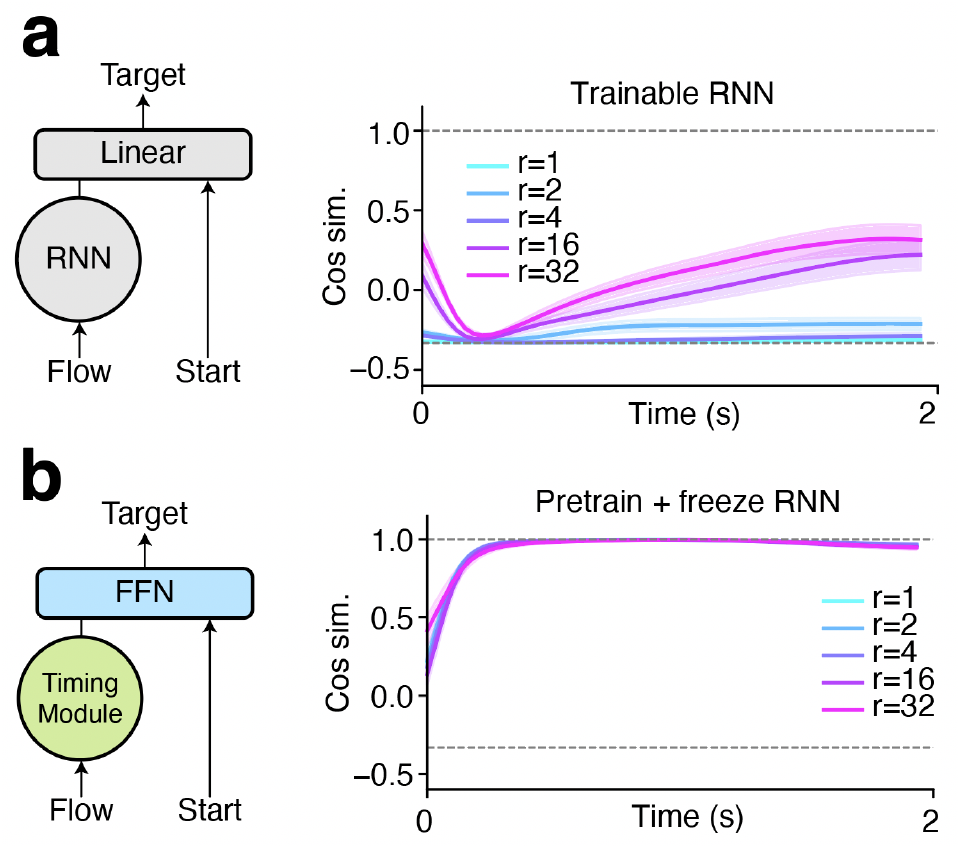
Pretrain-and-freeze, not rank, determines 1D integration. **a**, Unconstrained RNN + linear readout. An RNN with fully trainable recurrent weights receives Flow only; Start is concatenated at readout through a linear layer. Left, architecture schematic. Right, pairwise cosine similarity of velocity vectors across flow directions in the PCA space, swept over RNN rank (ranks 1, 2, 4, 16 and 32). All ranks remain below 1. **b**, Pretrained-and-frozen Timing Module + nonlinear FFN readout. The RNN is first trained to produce a monotonic timing ramp, then its recurrent weights are frozen; only the input weights and downstream FFN are trained on the task. Left, architecture schematic. Right, velocity cosine is near 1.0 for all ranks.

**Supplementary Figure S4.2.**
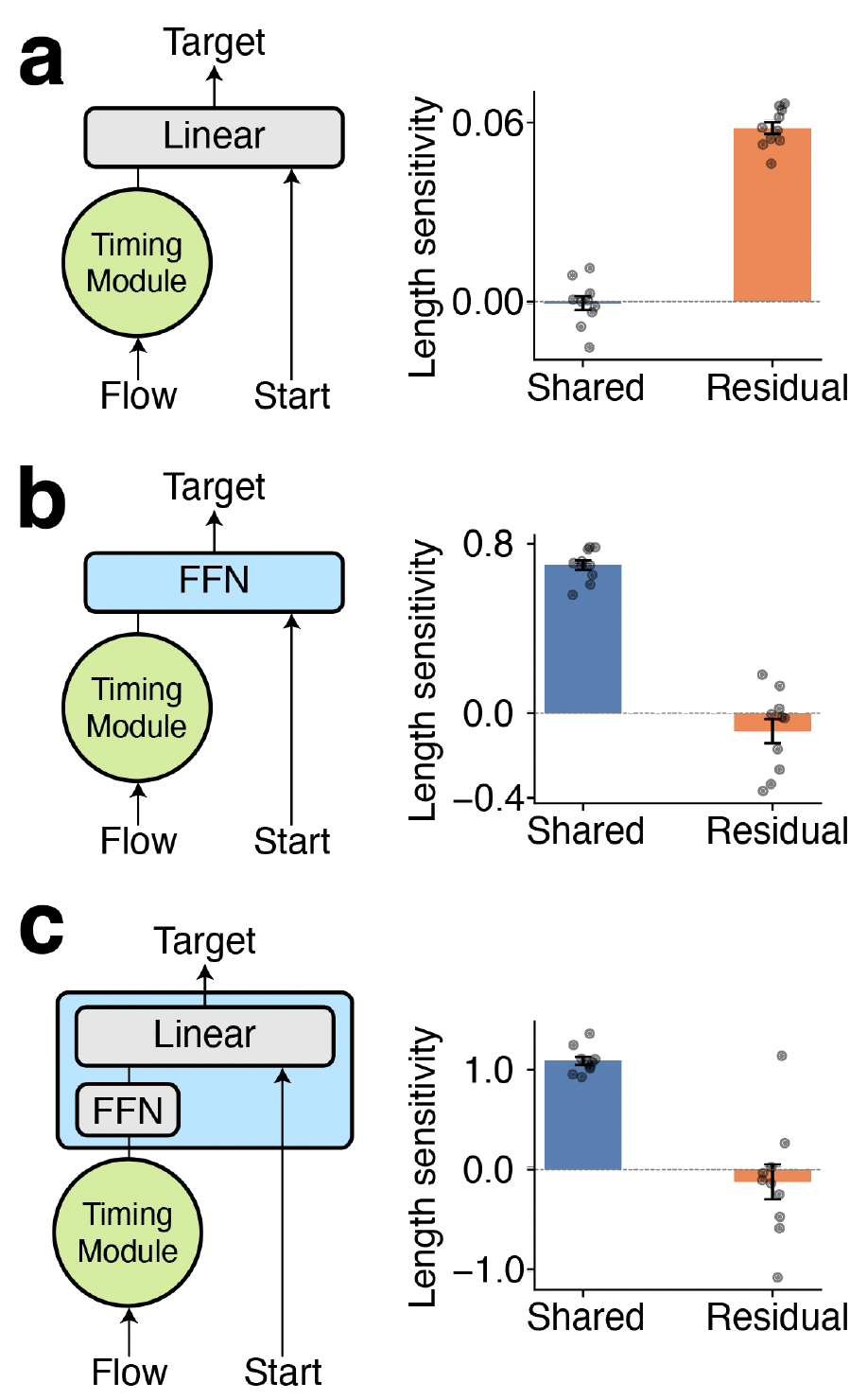
Causal perturbation analysis reveals how different readout architectures decode timing from the integrator. Two axes are defined from the integrator’s hidden-state trajectories for training conditions at magnitude 6. The common axis is the direction of the mean trajectory velocity averaged across all four cardinal flow directions. The residual axis is the deviation of a single direction’s endpoint from the mean endpoint; it captures direction-specific information caused by the recurrent nonlinearity. For each holdout condition, the hidden state h at flow offset is perturbed by ±ε (ε = 0.2) along each axis, the output displacement length is recomputed through the readout, and the signed slope (lengthh − length₋) / 2ε quantifies how sensitively the readout maps movement along that axis onto predicted magnitude. A positive slope indicates that perturbation along that axis consistently increases the predicted displacement length. Bars show mean ± s.e.m. across 10 seeds; individual seeds shown as dots. **a**, Timing Module + linear readout. The linear readout ignores the common axis (near-zero common slope) and instead reads length from the direction-specific residual (positive residual slope). **b**, Timing Module + FFN readout. The nonlinear readout extracts displacement length primarily from the common temporal axis (positive common slope, near-zero residual slope). **c**, Timing Module + task-aligned modular readout. Similar to b, the FFN sub-module in the readout extracts displacement length primarily from the common temporal axis.

**Supplementary Figure S4.3.**
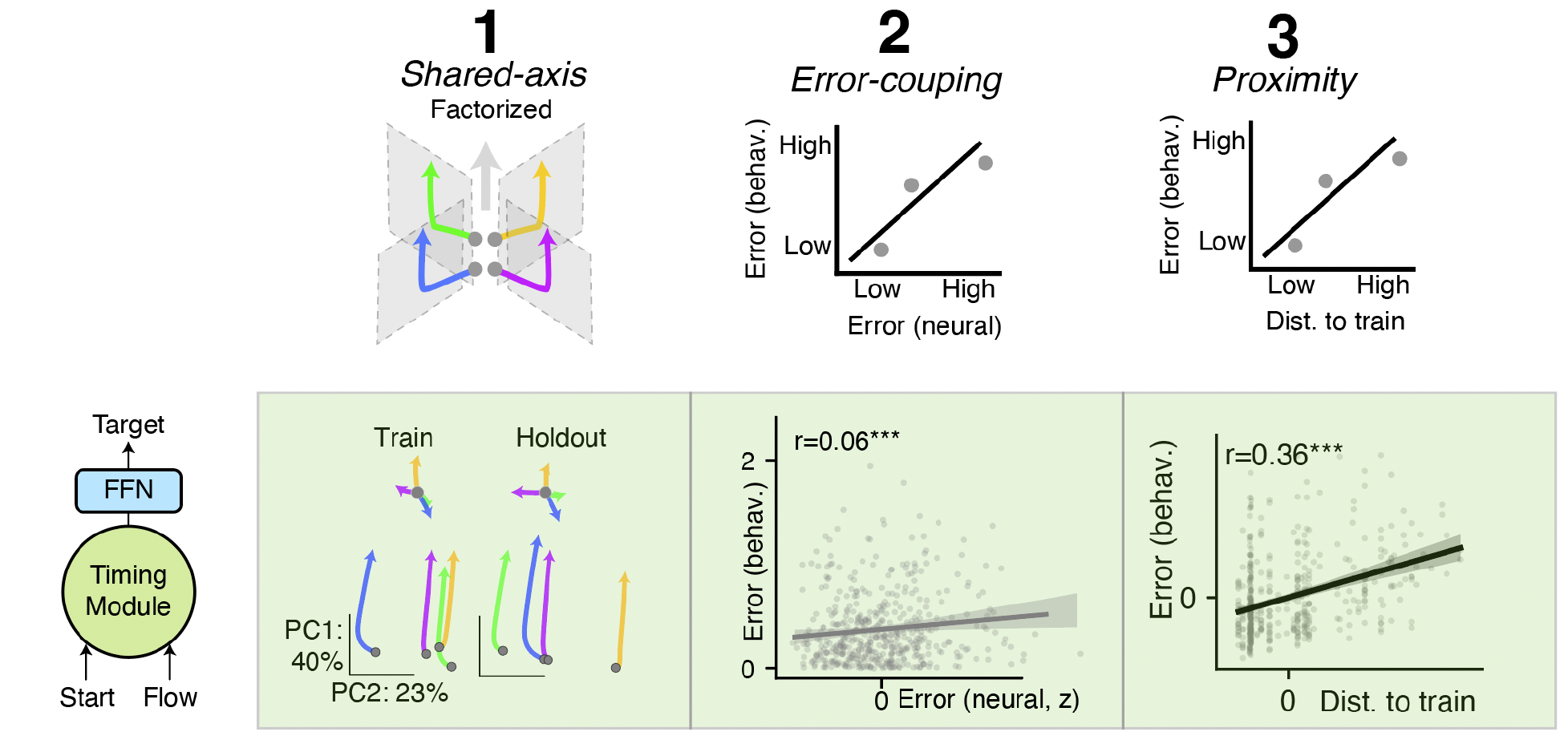
Additional control architecture. Timing Module with mixed input + FFN readout. Flow (4D) and Start (49D) are concatenated into a 53-dimensional input and fed jointly into the Timing Module, followed by the FFN transformation. Top: architecture schematic. Left, despite receiving both input streams, the Timing Module enforces a common temporal axis (PC1 = 40%), replicating 1D integration. Middle, neural deviation correlates with behavioral error (r = 0.06, p = 6.6 × 10⁻ ⁴). Right, proximity effect (r = 0.36, p = 1.5 × 10⁻¹ ⁶) emerges from the recurrent nonlinearity (tanh) that mixes Start and Flow at every timestep. This model replicates all three monkey features.

**Supplementary Figure S5.1.**
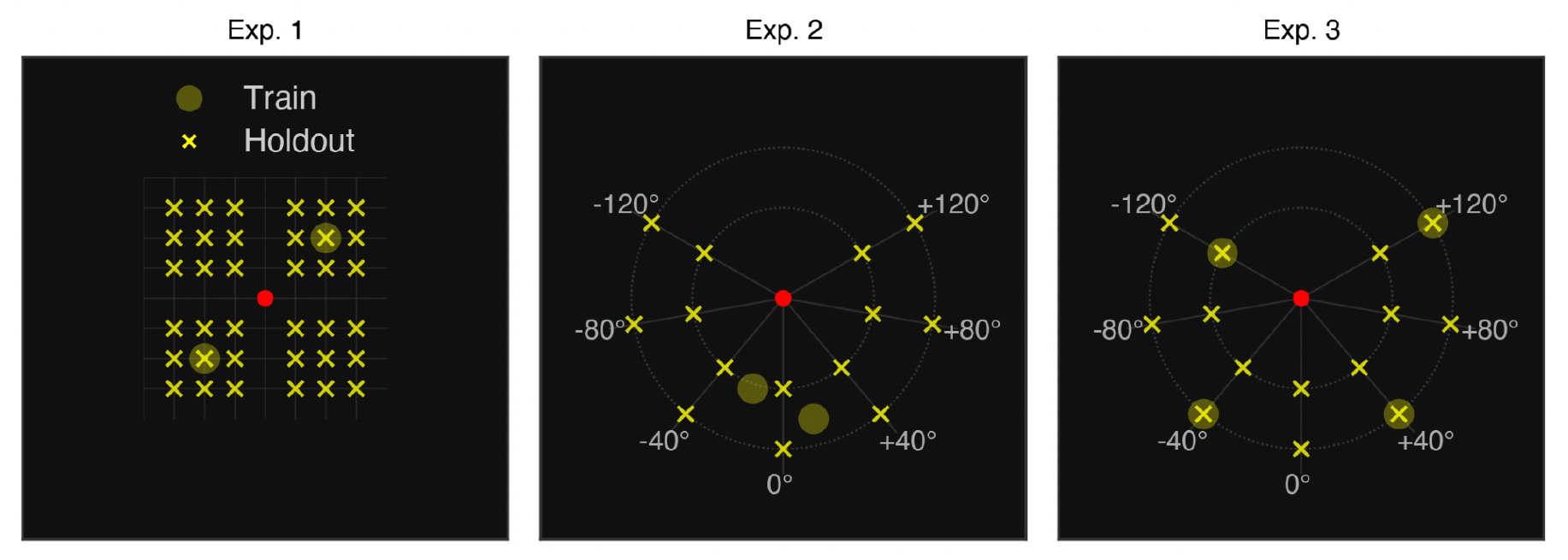
Training and holdout start positions of human psychophysics experiment. Each panel shows one experiment on the 800 × 800 px canvas. Yellow circles, training positions; yellow crosses, holdout positions; red dot, screen center. Dashed rings in Experiments 2 and 3 indicate the two concentric probe rings (radii 150 and 250 px). All dashed lines are not visible in the actual experiment. Angular labels denote deviation from the bottom of the screen (0° = directly below center). In Experiment 1, training positions are located at [−100, −100] and [+100, +100] relative to center, and holdout positions are distributed evenly across 36 locations on the canvas. In Experiment 2, training positions are concentrated near the bottom of the screen, where the Cartesian-to-polar mapping is maximally incompatible; holdout positions span ±120° from the bottom on both rings. Experiment 3 shares the same probe grid as Experiment 2; training positions are drawn from four angular lines (−120°, −40°, +40°, +120°), one ring per line.

**Supplementary Figure S5.2.**
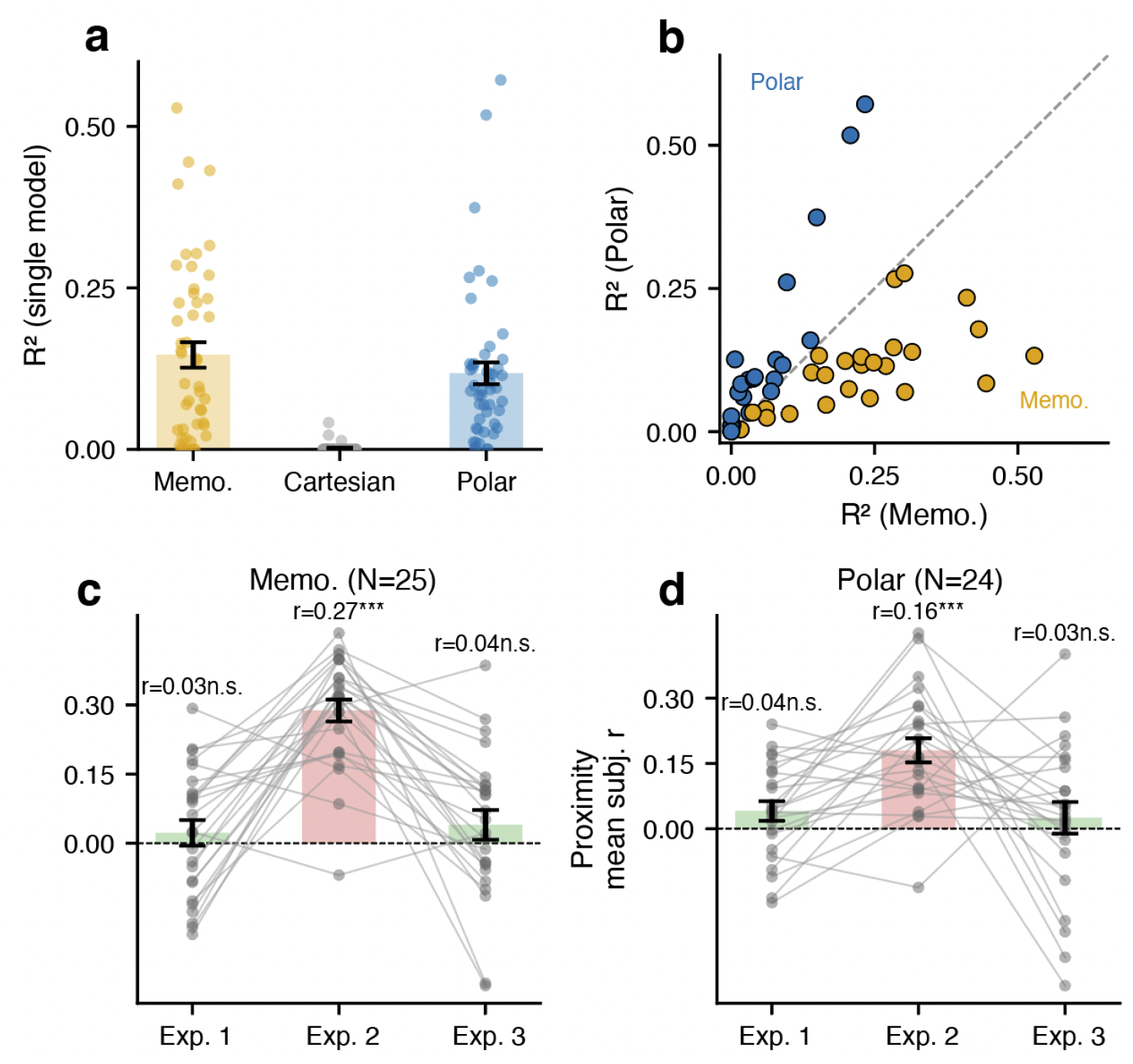
Strategy classification and proximity effect by group. **a**, We fitted each subject’s (*n* = 49) Experiment 2 holdout responses with three single-predictor models: a memorization model (mean displacement observed during training for the same flow direction, applied regardless of start position), a Cartesian model (Cartesian interpretation of the Flow), and a polar model (correct polar displacement from start to target). Bars show single-predictor R² (population means ± s.e.m.); each dot is one subject. Memorization and polar models both explained substantial variance among subjects, whereas the Cartesian model explained near-zero variance in all subjects. **b**, R²(memorization) versus R²(polar) for each subject. Subjects above the diagonal (blue) were classified as Polar (*n* = 24); subjects below (orange) as Memorization (*n* = 25). Dashed line, identity (classification boundary). **c–d**, Per-subject proximity correlation across the three experiments, shown separately for Memorization subjects (c, *n* = 25) and Polar subjects (d, *n* = 24). Proximity score is the Pearson r between distance to the nearest training condition (in 4D condition space) and normalized error of a holdout condition. Bars show group means ± s.e.m.; gray lines connect individual subjects across experiments. Both groups showed a significant proximity effect in Experiment 2 (Memorization: r = 0.27, p = 6.3 × 10⁻ ⁴ ²; Polar: r = 0.16, p = 1.7 × 10⁻¹ ⁵) but not in Experiments 1 or 3 (all n.s.). Memorization subjects produce a proximity gradient as expected, because the fixed trained displacement increasingly deviates from the correct polar target away from the training region. Polar subjects also showed the proximity effect, indicating that distance-dependent degradation is common for novel mappings regardless of the specific strategy adopted.

**Supplementary Figure S5.3.**
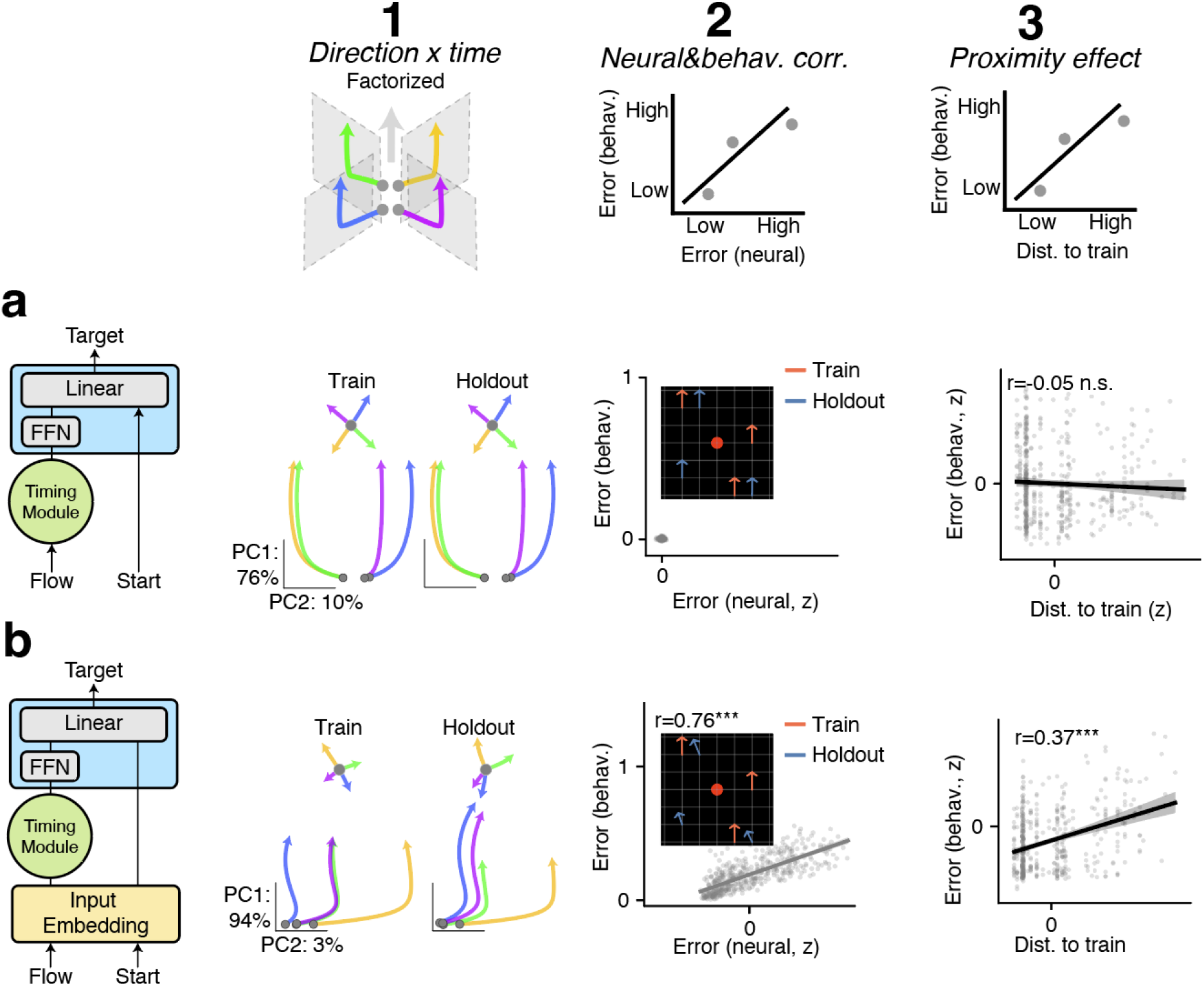
Model architectures for congruent and incongruent generalization. **a**, Full generalization model (Fig. 5f; Methods): an engineered architecture to replicate accurate compositional computation. Start and Flow enter through independent channels (factorized input). Flow is integrated by a dedicated Timing Module, and the readout is implemented by a modular decomposition: multiplication of direction × time (FFN) followed by addition with Start (linear layer). This model reproduced the shared integration axis observed in the monkey data and the accurate generalization observed in human Experiments 1 and 3: (1) one-dimensional shared integration (PC1 = 76%), (2) no spread of behavioral error across holdout conditions (fixed at 0), and (3) no proximity effect (r = −0.05, n.s.). **b**, Same architecture as the full generalization model, except that Start and Flow are jointly transformed by a nonlinear input embedding before entering the computation. This model replicated all three features observed in the monkey neural data and human behavior in Experiment 2: (1) one-dimensional shared integration (PC1 = 94%), (2) correlated neural deviation and behavioral error spanning a wide range (r = 0.76, p = 8.0 × 10⁻ ⁹ ⁰), and (3) significant proximity effect (r = 0.37, p = 1.9 × 10⁻¹ ⁷). The difference between this model and Model c (Fig. 4c) is that the readout implements a modular computation (displacement FFN + linear addition) rather than a non-specific FFN.

## Notes

### Competing Interest Statement

The authors have declared no competing interest.

